# Early life experience with natural odors modifies olfactory behavior through an associative process

**DOI:** 10.1101/2023.01.08.523155

**Authors:** Kristina V. Dylla, Thomas F. O’Connell, Elizabeth J. Hong

## Abstract

Past work has shown that chronic exposure of *Drosophila* to intense monomolecular odors in early life leads to homeostatic adaptation of olfactory neural responses and behavioral habituation to the familiar odor. Here, we found that, in contrast, persistent exposure to natural odors in early life increases behavioral attraction selectively to familiar odors. Odor experience increases the attractiveness of natural odors that are innately attractive and decreases the aversiveness of natural odors that are innately aversive. These changes in olfactory behavior are unlikely to arise from changes in the sensitivity of olfactory neurons at the first stages of olfactory processing: odor-evoked output from antennal lobe projection neurons was unchanged by chronic exposure to natural odors in terms of olfactory sensitivity, relational distances between odors, or response dynamics. We reveal a requirement for additional features of the environment beyond the odor in establishing odor experience-dependent behavioral plasticity. Passive odor exposure in a featureless environment lacking strong reinforcing cues was insufficient to elicit changes in olfactory preference; however, the same odor exposure resulted in behavioral plasticity when food was present in the environment. Together, these results indicate that behavioral plasticity elicited by persistent exposure to natural odors in early life is mediated by an associative process. In addition, they highlight the importance of using naturalistic odor stimuli for investigating olfactory function.

## INTRODUCTION

The structure and function of animal brains are shaped by sensory experience in early life. For instance, in the mammalian visual system, rearing animals in visual environments that contain only contours of a single orientation results in long-term changes in visual behaviors and selective shifts in cortical orientation maps to over-represent the experienced orientation^1–4^. Similarly, the spectrotemporal features of familiar sounds are over-represented and elicit stronger responses in rodent cortical auditory neurons^5,6^. In the olfactory system, odor experience in early life also impacts how animals behave towards familiar odors later in life. For instance, divergent olfactory experiences arising from childhoods in different cultures or from prenatal exposure to different maternal diets can alter the perception of odor intensity or pleasantness ^7–11^. In vertebrate and invertebrate animals, persistent passive exposure to odors in early life elicits long-lasting changes in the structure and function of olfactory systems and in odor-dependent behaviors^12,13^.

One hypothesis is that experience-dependent plasticity helps sensory systems adapt to the statistical distribution of stimuli in the local environment, promoting the efficient encoding of sensory information with a limited number of neurons^14–18^. However, raising animals in highly controlled environments dominated by a single defined stimulus (e.g., contour orientation, sound frequency, monomolecular odorant), often degrades performance in discrimination tasks that depend on the familiar stimulus and can have a deleterious impact on sensory tasks that use unfamiliar stimuli^3,19,20^. Thus, laboratory studies using extreme artificial sensory environments provide insights into the instructive roles of normal sensory experience in the developing nervous system but may also reflect the consequences of prolonged exposure to abnormal sensory environments.

Broadly, the goal of this study was to better understand how sensory experience in early life affects sensory coding and behavior in natural environments. We investigated this problem in the olfactory system of the vinegar fly *Drosophila melanogaster*, a well characterized sensory system with several experimental advantages, which include its numerical compactness, genetic accessibility, and the availability of powerful tools for functional circuit interrogation^21,22^. In *Drosophila*, chronic exposure of flies to high concentrations of a monomolecular odor reduces behavioral responses towards that odor^13,17,18,23,24^. This effect is selective for the specific odor that is common or overrepresented in the environment; responses to novel unrelated odors are unaffected. Behavioral plasticity triggered by chronic exposure to high concentration of monomolecular odor has been attributed to long-term, stimulus-specific increases in inhibitory gain at the first synaptic stage of olfactory processing^17,18^. Very high concentrations of monomolecular odors, as were used in these studies, are nearly always innately aversive to animals, and long-term behavioral habituation to such odors would be beneficial by allowing flies to occupy environments naïve animals find aversive.

Monomolecular odor stimuli used in prior studies of olfactory plasticity diverge from natural odors in several important ways. First, natural odors are complex blends of many distinct monomolecular odorants, often mixed at characteristic ratios^25–27^. Second, individual monomolecular volatiles in natural odor stimuli are present at significantly lower concentrations^28–30^ (∼1 ppb to <10 ppm in air) compared to monomolecular concentrations presented in prior laboratory studies (∼10^3^–10^4^ ppm)^17,18,23^. Finally, many natural odors elicit behavioral attraction^31,32^, even at the highest naturally occurring intensities that would be encountered close to the odor source.

Here, we asked whether long-lasting experience with ecologically relevant odors in early life, particularly innately attractive odors such as those arising from appetitive natural food sources, also elicits olfactory plasticity in flies, and, if so, what form that plasticity takes. The answer is not straightforward since, if behavioral plasticity induced by chronic odor exposure is primarily driven by sensory habituation and decreased gain at the first olfactory synapse, flies should respond less strongly (be less sensitive) to the familiar odor. However, from the perspective of what is adaptive for survival, one might predict that flies should respond more strongly (be more sensitive) to a familiar odor from an environment that supported survival and reproduction in early life.

We investigated this question by rearing animals in environments odorized with natural food sources that could be realistically encountered in the biological world. We found that chronic exposure to an innately attractive odor can increase behavioral attraction selectively towards that odor, and exposure to an innately aversive odor can decrease behavioral aversion towards that odor. Unexpectedly, functional imaging studies showed that olfactory projection neuron output is stable and unaffected by chronic exposure to odors in naturalistic conditions, indicating changes in olfactory gain or sensitivity are unlikely to account for behavioral plasticity. Using a novel apparatus for rearing flies in odorized environments, we demonstrate that access to food during odor experience is required to elicit behavioral plasticity. Together these results demonstrate that changes in olfactory behavior elicited by long-term experience with natural odors in early life occurs through a non-classical form of associative learning.

## Results

### Evaluating the attractiveness or aversiveness of natural odors

We used a free-flight trap-based assay to evaluate the behavioral attractiveness or aversiveness of odors arising from various natural food sources. Trap-based assays were chosen because they naturalistically model odor-guided food search by hungry flies; video monitoring of the flies’ behavior near the trap entrances allowed us to survey different stages of fly foraging behavior (Figure 1A). Behavioral attraction was measured by comparing the number of flies that entered a trap baited with the odor source to the number entering a control trap containing water (Figure 1Ai). The top of each trap had five openings, each ∼1 mm in diameter, through which odor diffused out of the trap and through which flies could enter into the trap. The trap openings were centered in a 10-cm circular platform on which flies could land and explore the openings. Each trap was simultaneously imaged from the top and side view (Figure 1Aii-iii).

**Figure 1:**
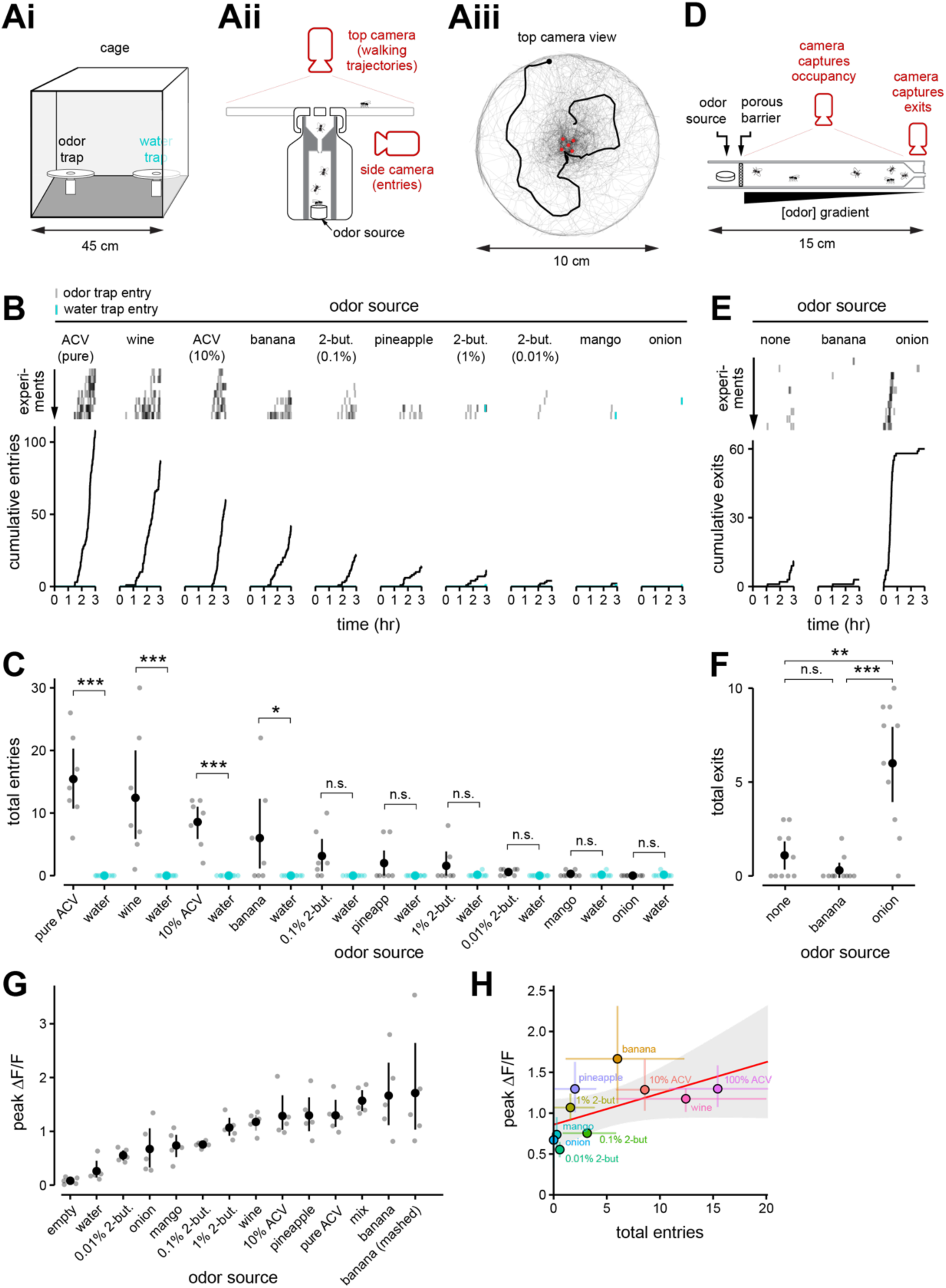
Natural odor sources vary in their innate behavioral attractiveness to flies in a free flight trap-based olfactory assay. **(Ai - Aiii)** Schematics of free-flight trap-based assay. **(Ai)** Flies are presented with two traps, one containing an odor source and one containing water, within a mesh-enclosed cubic cage. **(Aii)** Cross-section through an odor trap. The top camera records the flies’ walking behavior on the trap platform, surrounding the trap entrances. The side camera records the flies’ passages into the trap. Flies are unable to exit the trap once they have passed through the narrow entrance passage. (Aiii) Example reconstructed walking trajectories (gray traces) on the platform of a trap baited with banana. One individual trajectory is shown in black, with the endpoint designated with a black dot. Red circles mark the entrance apertures of the trap. **(B)** Cumulative trap entries for ten different odor sources, each tested against water. Groups of fifty starved flies choose between a trap baited with an odor source (black) and a trap containing water (blue). Top: Time of individual entries in each experiment (row) for each odor source, experiments sorted by the total number of entries. Bottom: Cumulative entries across all experiments (n = 7 experiments; ACV, apple cider vinegar; 2-but., 2-butanone). **(C)** Mean and 95% CI for total entries from (**B)** into traps baited with each odor source (black) tested against water (blue) (n = 7 experiments). \**p* < 0.05, \*\**p* < 0.01, \*\*\**p* < 0.001, n.s., no significant difference (*p* ≥ 0.05), two-way analysis of variance with Bonferroni correction for multiple comparisons. **(D)** Cross-sectional view of the exit assay. Groups of ten unstarved flies are introduced into the main chamber, constructed of transparent plastic. A porous barrier prevented flies from physically touching the odor source but allowed odor to diffuse into the main chamber. The other side of the tube narrowed to a small opening through which flies could exit. Clean air diffused into the chamber through the exit port, establishing an odor gradient between the odor source and the exit. The flies’ positions and exits from the trap were recorded for three hours. **(E)** Cumulative exits from the exit assay. Three odor sources were individually tested in parallel: banana, onion, or no odor (control). Top: Time of individual exits in each experiment (row) for each odor source, sorted by the time of first exit. Bottom: Cumulative exits across all experiments (n = 10). **(F)** Mean and 95% CI for total exits from (**E)** from traps baited with each odor source (n = 10 experiments). \*\**p* < 0.01, \*\*\**p* < 0.001, n.s., no significant difference (*p* ≥ 0.05), two-tailed permutation t-test with Bonferroni correction. **(G)** Peak amplitude (mean and 95% CI, solid symbol) of stimulus-evoked calcium responses in projection neuron (PN) terminals, measured in a single plane of the mushroom body (MB) calyx in flies expressing GCaMP6f in ∼75% of PNs (n = 5 flies; light symbols are average of three trials per stimulus in each fly). ΔF/F response was computed as the pixel mean within a large ROI that circumscribed PN axonal boutons in each fly. See Supplementary Figure S1C for mean response traces (changes in fluorescence over time). Odor stimuli from left to right: empty odor vial; water; 2-butanone (10^-4^); onion; mango; 2-butanone (10^-3^); 2-butanone (10^-2^); red wine; 10% apple cider vinegar (ACV); pineapple; 100% ACV; mix of 2-butanone, 2-pentanone, E2-hexenal, and pentyl acetate, each at 1%; banana; and mashed banana. The fly genotype was *yw/+; NP225-Gal4/+; 20x-UAS-IVS-Syn21-OpGCaMP6f-p10/+; +*. **(H)** Peak amplitudes of odor-evoked PN responses (mean and 95% CI, from **(G)**) are weakly correlated with total trap entries (mean and 95% CI) elicited by each odor from **(C)**; Spearman’s rank-order correlation, *r* = 0.76, *p* = 0.016.

In each experiment, fifty flies (3-4 days old; starved for 24 hr) were introduced into a cage (∼90 dm^3^) containing an odorized trap and a water trap, and their positions and entries at the traps were measured for three hours. Entries of individual flies into each trap were extracted from side video data (see Methods) and revealed that the odors of the ten sources tested varied in their behavioral attractiveness (Figure 1B). The odors of apple cider vinegar and red wine were the most attractive in our stimulus panel, confirming prior reports of their reliability as olfactory attractants^33–35^, and the odor of banana fruit was moderately attractive (Figure 1B-C). The behavioral attractiveness of odors was concentration-dependent: flies were more attracted to the odor of undiluted apple cider vinegar compared to 10% apple cider, and they were most attracted to the odor of 0.1% 2-butanone compared to either 1% or 0.01% dilutions. Entries into the water trap in any experiment with any odor source were rare (Figure 1C), and many flies do not enter either trap within the three hours assayed.

Entries into the onion trap were not observed in any of ten replicate experiments. To determine whether flies avoid the odor of onion or, alternatively, either do not detect the odor of onion or behave neutrally towards it, we developed an “exit assay” that measures odor avoidance as the number of flies that leave an odorized environment compared to an unodorized environment (Figure 1D). Groups of ten unstarved flies were introduced into a cylindrical chamber (25 cm^3^) odorized with banana, onion, or no odor (control). Whereas flies typically exited the odorized environment within the first hour of the three-hour experiment when onion was the odor source (Figure 1E), significantly fewer flies left when it was not odorized (control) or odorized with banana (Figure 1F). These results indicate that flies detect and avoid the odor of onion in this assay.

To evaluate the relationship between the attractiveness of an odor and the sensitivity of the fly olfactory system for the odor, we estimated the overall level of olfactory activity elicited by each odor using two-photon functional imaging. We expressed the genetically encoded calcium indicator GCaMP6f in ∼75% of olfactory projection neurons (PNs; from the *NP0225-GAL4* driver^36^) and measured odor-evoked calcium signals from PN axon terminals where they arborize in the calyx of the mushroom body (Figure S1A). We confirmed that the relative amplitudes of bulk PN bouton responses among stimuli is not strongly sensitive to the specific imaging plane in the calyx (Figure S1A-B). All odor stimuli in our panel evoked reliable, widespread calcium signals in PN terminals throughout the calyx (Figure 1G, S1A-C). Additionally, odors that were more behaviorally attractive tended to evoke higher mean levels of olfactory activity than odors that were less attractive (Spearman’s rank correlation, *r*=0.76, *p*=0.016) (Figure 1G-H, Figure S1C). These results confirm that the odor stimuli used in our behavioral experiments are detected by the fly olfactory system and show that the level of olfactory activity elicited by an odor is related to how attractive it will be to naïve flies.

### Chronic exposure to a natural odor in early life increases behavioral attraction selectively towards that odor

We asked whether chronic exposure of flies to odors from natural sources in early life can change the attractiveness or aversiveness of the odor when it is encountered again later in life. We continually exposed groups of newly eclosed flies for two days to the odor of banana (moderately attractive), onion (aversive), or to no odor (control; Figure 2A). Flies could not physically contact the odor sources, and odor exposure itself did not impact the survival of flies (Figure 2B). Following odor exposure, flies were wet-starved for 24 hours in the absence of the odor and then tested for three hours in the trap assay (Figure 1A). In these experiments, flies were presented with a choice between a trap baited with banana and a trap baited with onion (Figure 2C).

**Figure 2:**
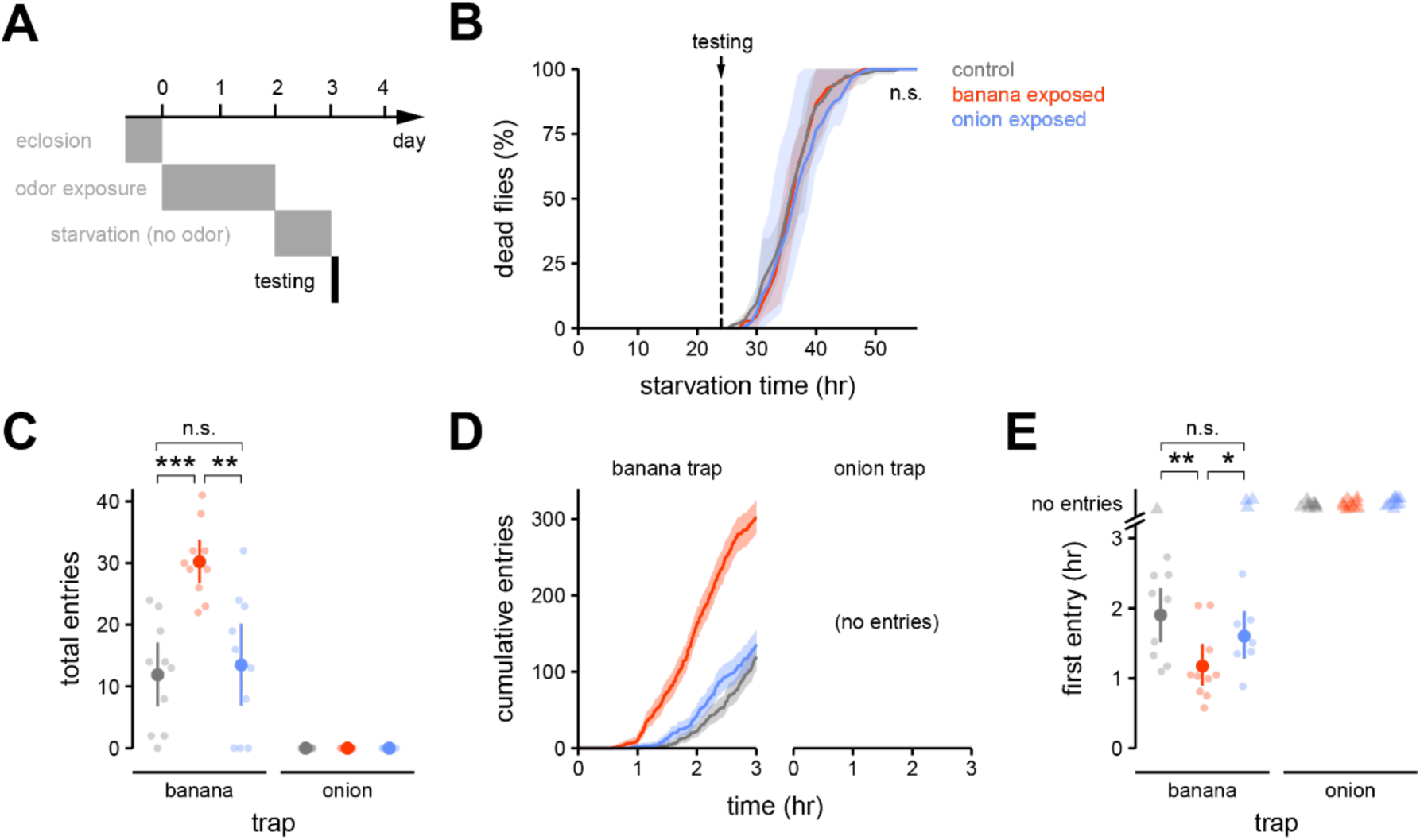
Persistent exposure to banana odor in early life increases behavioral attraction selectively to banana odor. **(A)** Newly eclosed flies were exposed to a natural odor source for two days, then starved for one day on wet tissue paper in the absence of the odor source. Flies were presented with a choice between a trap baited with onion or a trap baited with banana (Figure 1A) for the duration of three hours. Olfactory preference was evaluated in 3–4 day old flies. **(B)** Survival for control- (grey), banana odor- (red), or onion odor- (blue) exposed flies (mean and 95% CI; n = 3 experiments/group) as a function of starvation time. Odor exposure as in **(A)**; starvation time 0 corresponds to the start of the third day in **(A).** Testing would normally commence at 24 hr starvation. Odor experience does not affect survival, n.s., no significant difference, *p* ≥ 0.05 (two-sample Kolmogorov-Smirnov test with Bonferroni correction). **(C)** Total entries (mean and 95% CI, solid symbols) into the banana- or onion-odorized trap over three hours by control- (grey), banana odor- (red), or onion odor- (blue) exposed flies (n = 10 experiments, light symbols). No flies entered the onion trap. \*\**p* < 0.01, \*\*\**p* < 0.001, n.s., no significant difference (*p* ≥ 0.05), two-tailed permutation t-test with Bonferroni correction. **(D)** Cumulative entries into each trap across all experiments for each odor exposure group from **(C)**. Error envelope is the 95% CI bootstrapped across experiments (n = 10). **(E)** Time of the first entry into each trap in an experiment (mean and 95% CI, solid symbol) from **(C)** (individual experiments, light symbol). Triangle symbols mark cases where no entries were observed. \**p* < 0.05, \*\**p* < 0.01, n.s., no significant difference (*p* ≥ 0.05), pairwise log-rank test with Bonferroni correction.

Of the 500 flies assayed across ten experiments, none were observed to enter the onion trap (Figure 2C-D). Thus, prior exposure to onion odor was unable to convert the innate aversion of flies towards onion odor into attraction. However, chronic exposure of flies to the odor of banana resulted in a stronger preference for entering the banana-baited trap, as compared to control-exposed flies. Exposure to banana odor increased the mean number of total entries into the banana trap (Figure 2C) and also decreased the mean latency to the first entry of a fly into the banana trap (Figure 2E). This effect was odor-specific, since onion odor-exposed flies entered the banana trap at similar rates as controls (Figure 2C-D). Thus, chronic exposure of flies to odors from natural sources can cause a long-lasting change in olfactory behavior, and this change is observed selectively only for the familiar odor.

We wondered whether the earlier and more frequent entry of banana odor-exposed flies into the banana trap reflected a heightened ability to detect and/or navigate to the banana odor. To examine different stages of the flies’ odor-guided search, particularly near the trap entrances where the odor concentration is highest, we extracted the walking trajectories of individual flies on the platforms of the banana or onion traps, using data from the top view camera (Figure 3A, S2A). These data showed that flies from all odor exposure groups landed at least briefly on both odor traps (Figure 3B-C, S2B), but they occupied and explored the surface of the banana trap much more extensively throughout the experiment (Figure 3B-E, S2A). Focusing on the banana trap, we observed that, as a population, flies from all odor exposure groups located and landed on the banana trap platform with similar latencies (Figure 3B-C, S2B) and spent a similar amount of time on the trap platform during the first 30 min of each experiment (Figure 3D-E). These observations suggest that odor experience does not strongly affect the behavioral sensitivity or ability of the flies to locate the familiar odor in this assay.

**Figure 3:**
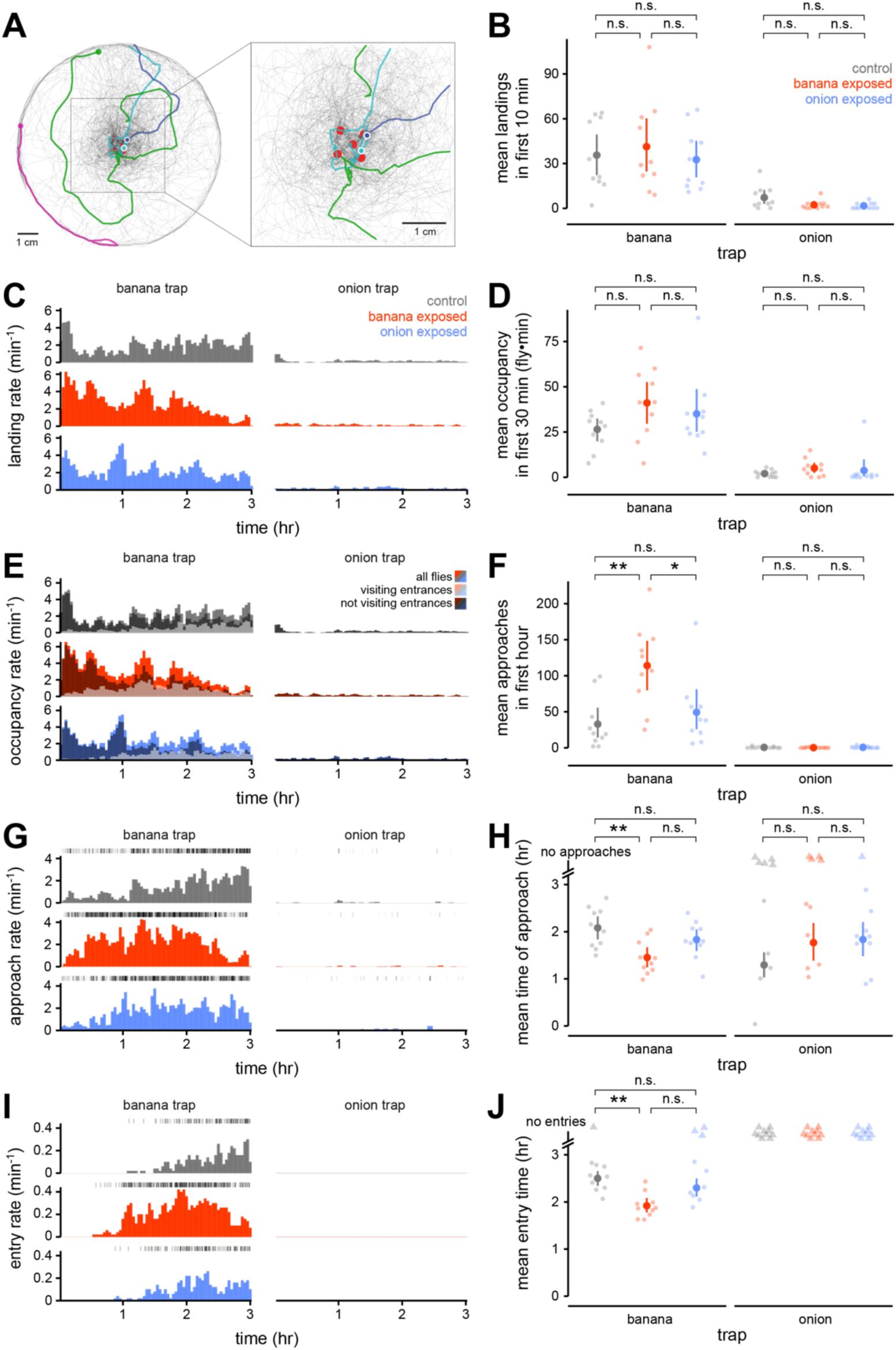
Chronic olfactory experience alters different stages of odor-guided search behavior towards the familiar odor. **(A)** Example reconstructed walking trajectories (light grey traces) on the platform of a trap baited with banana with four individual trajectories highlighted in color. The solid circle marks the trajectory endpoint. The five red circles comprise the region of interest (ROI) containing the entrance holes to the trap. Inset: enlarged view of the trap entries. The cyan, and blue traces are “approaches,” defined as trajectories that enter the ROI at least once, whereas the green and magenta traces are not. **(B - J)** Quantification of fly behavior in experiments from Figure 2C-E. Control- (grey), banana odor- (red), or onion odor- (blue) exposed flies in the free-flight assay chose between a trap baited with onion or a trap baited with banana (n = 10 experiments/condition). For **B**, **D**, **F**, **H**, **J**, solid symbols are mean and 95% CI, shaded symbols are individual experiments. \**p* < 0.05, \*\**p* < 0.01, n.s., no significant difference (*p* ≥ 0.05), two-tailed permutation t-test with Bonferroni correction. For **C**, **E**, **G**, **I**, histograms report the event rate computed in 5-min bins with 50% overlap. **(B)** Mean number of landings on each trap during the first ten minutes of each experiment. **(C)** Landing rate over time across all experiments in each condition. Landings are defined as the initiation of a trajectory. **(D)** Mean occupancy on each trap during the first 30 minutes of each experiment. Occupancy was measured in units of fly•min; one fly on the trap for 5 minutes is equivalent to five flies on the trap for 1 minute each. **(E)** Occupancy rate over time across all experiments in each condition. Occupancy was measured as the number of unique trajectories occurring in the time bin. Occupancy is broken down into flies that do not visit the trap entries (trajectories out of ROI, dark shaded) and flies that visit the trap entries (pass through the ROI at least once, light shaded) during each time bin. **(F)** Mean number of approaches (to entrances) on each trap during the first hour of the experiment. **(G)** Approach rate (to entrances) over time across all experiments in each condition. Raster plots above each histogram show the time of each approach. **(H)** Mean (across experiments) of the median time of approach in each experiment for each condition. **(I)** Entry rate over time across all experiments in each condition. Raster plots above each histogram show the time of each entry. No entries into the onion trap were observed in any experiments. **(J)** Mean (across experiments) of the median time of entry in each experiment for each condition.

Comparisons of occupancy on the trap platform measured from the top view camera (Figure 3C, E) with trap entries measured with the side view camera (Figure 2D) showed that flies from all odor exposure conditions locate the banana trap much earlier than when they enter it (Figure 3C, I). This observation suggests that flies evaluate the trap for some time before making the decision to enter. We examined movies of the flies’ behavior at the entrances of the banana trap and found that flies rarely enter the trap on their first visit to an entrance. They typically investigated one or more entrance holes multiple times before entering the trap, often repeatedly extending and retracting their head and upper thorax into the apertures (Supplemental Video S1). We defined a region of interest (ROI) that comprised the (noncontiguous) set of pixels containing the five trap entrances (Figure 3A, red), approximately 1% of the total area of the platform. On average, as the experiment progressed, the fraction of flies on the trap platform that visited the ROI (i.e., at the trap entrances) increased relative to the fraction not visiting the ROI (Figure 3E), consistent with an increasing intensity of investigation at the source of the odor. We defined a walking trajectory in which flies entered into any part of the ROI as an “approach” to the trap entrances (Figure 3A, inset, for approach examples) and compared their frequency and timing to that of entries into the trap (Figure 3G, I).

For all groups of flies, the number of approaches to the trap entrances was much higher than the number of entries into the banana trap, and entries into the banana trap began much later than approaches (Figure 3G, I). Compared with onion odor- or control-exposed flies, banana odor-exposed flies approached the entrances of the banana trap earlier (median time, 1.45 hr for banana odor-exposed versus 1.83 hr or 2.08 hr for onion odor- or control-exposed, respectively) (Figure 3G-H) and more frequently (Figure 3F-G). Other metrics of fly behavior, including walking speeds (Figure S2A, S2C, S2E), spatial distribution on the trap (Figure S2D), and time duration of trajectories (Figure S2F-G), were unaffected by chronic odor exposure. Thus, olfactory experience does not appear to impact the ability of flies to locate the banana trap but affects odor-guided behavior at the trap in this assay. Flies more readily and more vigorously investigate a local source of the familiar odor when they re-encounter it, concomitant with earlier and more frequent entries into the trap baited with the experienced odor. These results are most consistent with chronic experience-dependent behavioral plasticity representing a change in the meaning or mapping of the experienced odor to behavioral programs for approach, rather than a change in the sensitivity of the fly to the odor.

### Olfactory experience can alter the naïve rank preference for different natural odors

We tested whether the form of behavioral plasticity elicited by chronic exposure to banana odor also extends to additional odors and additional choice contexts. When given a choice between two traps, one odorized with pineapple and one containing water, flies exposed to the odor of pineapple entered traps odorized with pineapple earlier and in larger numbers, compared to flies exposed to banana odor or no odor (control) (Figure 4A-B). The latency to the first entry into the pineapple trap tended to be earlier in the pineapple odor-exposed condition across experiments, though the trend was not statistically significant (Figure 4C). Flies in the banana odor-exposed and control conditions were statistically indistinguishable across all metrics, confirming that behavior plasticity was odor-specific. These results demonstrate that heightened behavioral attraction elicited by prior experience with attractive natural odors generalizes to another attractive odor and occurs regardless of whether flies are choosing between attractive and aversive stimuli (banana vs. onion odor) or attractive and neutral stimuli (pineapple vs. water odor).

**Figure 4:**
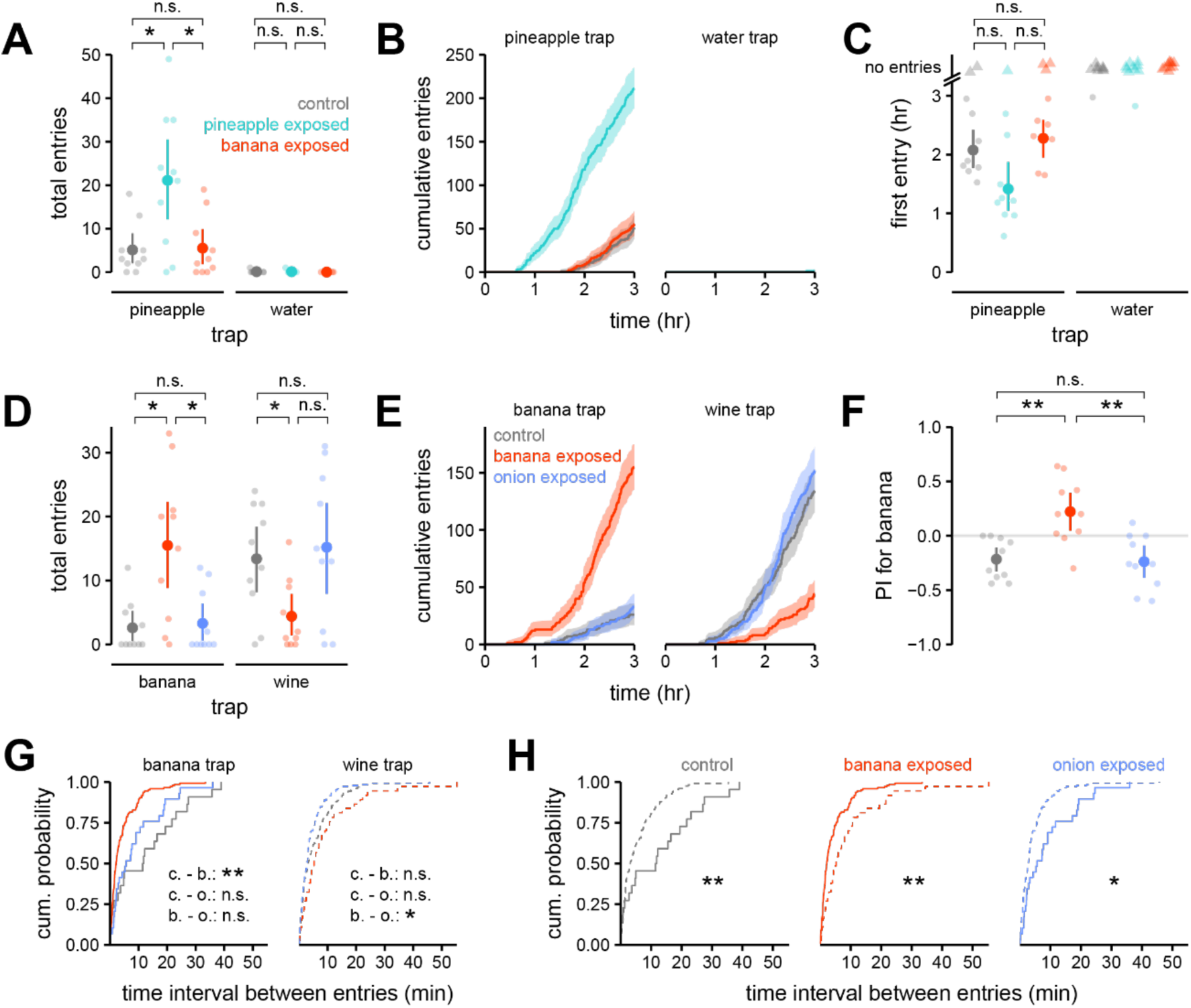
Behavioral plasticity elicited by odor experience in early life extends to additional odors and additional choice contexts. **(A)** Total entries (mean and 95% CI, solid symbols) over three hours into the pineapple- or water- (control) odorized trap by control- (gray), banana odor- (red), or pineapple odor- (cyan) exposed flies (n = 10 experiments, light symbols). Free flight trap-based assay was as in Figure 2, except that groups of seventy, rather than fifty, flies were tested in each experiment. \**p* < 0.05, n.s., no significant difference (*p* ≥ 0.05), two-tailed permutation t-test with Bonferroni correction. **(B)** Cumulative entries into each trap across all experiments (n = 10) for each condition from **(A)**. Error envelope is the 95% CI bootstrapped across experiments. **(C)** Time of first entry (mean and 95% CI, solid symbol) into each trap for each experiment (light symbols) in **(A)**. Triangle symbols denote experiments where no entries were observed. n.s., no significant difference (*p* ≥ 0.05), pairwise log-rank test with Bonferroni correction. **(D)** Total entries (mean and 95% CI, solid symbols) over three hours into the banana- or wine-odorized trap by control- (gray), banana odor- (red), or onion odor- (blue) exposed flies (n = 10 experiments, light symbols). \**p* < 0.05, n.s., no significant difference (*p* ≥ 0.05), two-tailed permutation t-test with Bonferroni correction. **(E)** Cumulative entries into each trap across all experiments (n = 10) for each condition from **(D)**. Error envelope is the 95% CI bootstrapped across experiments. **(F)** Preference index for banana (mean and 95% CI, solid symbols) for each odor exposure group from **(D)**. \*\**p* < 0.01, n.s., no significant difference (*p* ≥ 0.05), two-tailed permutation t-test with Bonferroni correction. **(G-H)** Cumulative probability distributions of inter-entry time intervals for experiments in **(D)**, grouped by odor trap **(G)** or odor exposure condition **(H).** Solid curves, banana trap; dashed curves, wine trap. \*\**p* < 0.01, \**p* < 0.05, n.s., no significant difference (*p* ≥ 0.05), two-sample Kolmogorov-Smirnov test with Bonferroni correction.

We next asked whether olfactory experience can alter the naïve relative preference of flies for different odors. Naïve (and control-exposed) flies reliably preferred the odor of wine over that of banana in the trap assay (Figure 1B-C, Figure 4D). However, when given a choice between wine- and banana-odorized traps, banana odor-exposed flies entered the banana trap earlier and in greater numbers compared to flies exposed to no odor (control-exposed) (Figure 4D-F). The change in preference was odor-specific; onion odor-exposed flies and control flies exhibited similarly strong preferences for wine odor over banana odor (Figure 4F). Banana odor-exposed flies entered the banana trap at a higher frequency than the wine trap, whereas the opposite was true for onion odor-exposed and control flies (Figure 4G-H). Therefore, the increased number of entries of banana odor-exposed flies into the banana trap during the three-hour testing period was attributable not only to an earlier onset of entries (as in Figure 2D), but also a higher rate of entries. These data show that prior experience with a natural odor in early life can increase its attractiveness sufficiently to alter the relative preference of the fly for that odor in relation to another attractive stimulus.

### Persistent experience with an aversive natural odor in early life reduces behavioral avoidance of the familiar odor

Exposure in early life to the odor of onion, which is aversive to naïve flies, did not significantly alter how flies behaved towards any odor, including onion odor, in the free flight trap-based assay (Figure 1B-C). This result might indicate that chronic exposure to onion odor does not alter fly olfactory behavior; however, another possibility is that, because the trap assay mostly measures behavioral attraction, it is poorly suited to reporting changes in the behavioral aversiveness of an odor.

To evaluate behavioral aversion more directly, we tested the olfactory behavior of unstarved control and onion odor-exposed flies using the exit assay (Figure 5A; Figure 1D-F). We measured the flies’ positions in, and exits from, three different environments, each odorized with onion, banana, or no odor source. The mean number of total exits during the three-hour testing period was not significantly different between odor exposure groups from all odor environments (Figure 5B), but the cumulative number of exits (across experiments) from the onion-odorized environment was ∼30% reduced in onion odor-exposed flies compared to controls (Figure 5C). Additionally, across all flies in all experiments for each exposure group, the median time spent in the onion-odorized environment was higher for onion odor-exposed flies as compared to control flies (Figure 5D), consistent with fewer exits among onion odor-exposed flies. This difference in dwell times was observed in the onion-odorized environment, but not the banana-odorized or non-odorized environments, indicating that the change in olfactory behavior was specific to the exposure odor.

**Figure 5:**
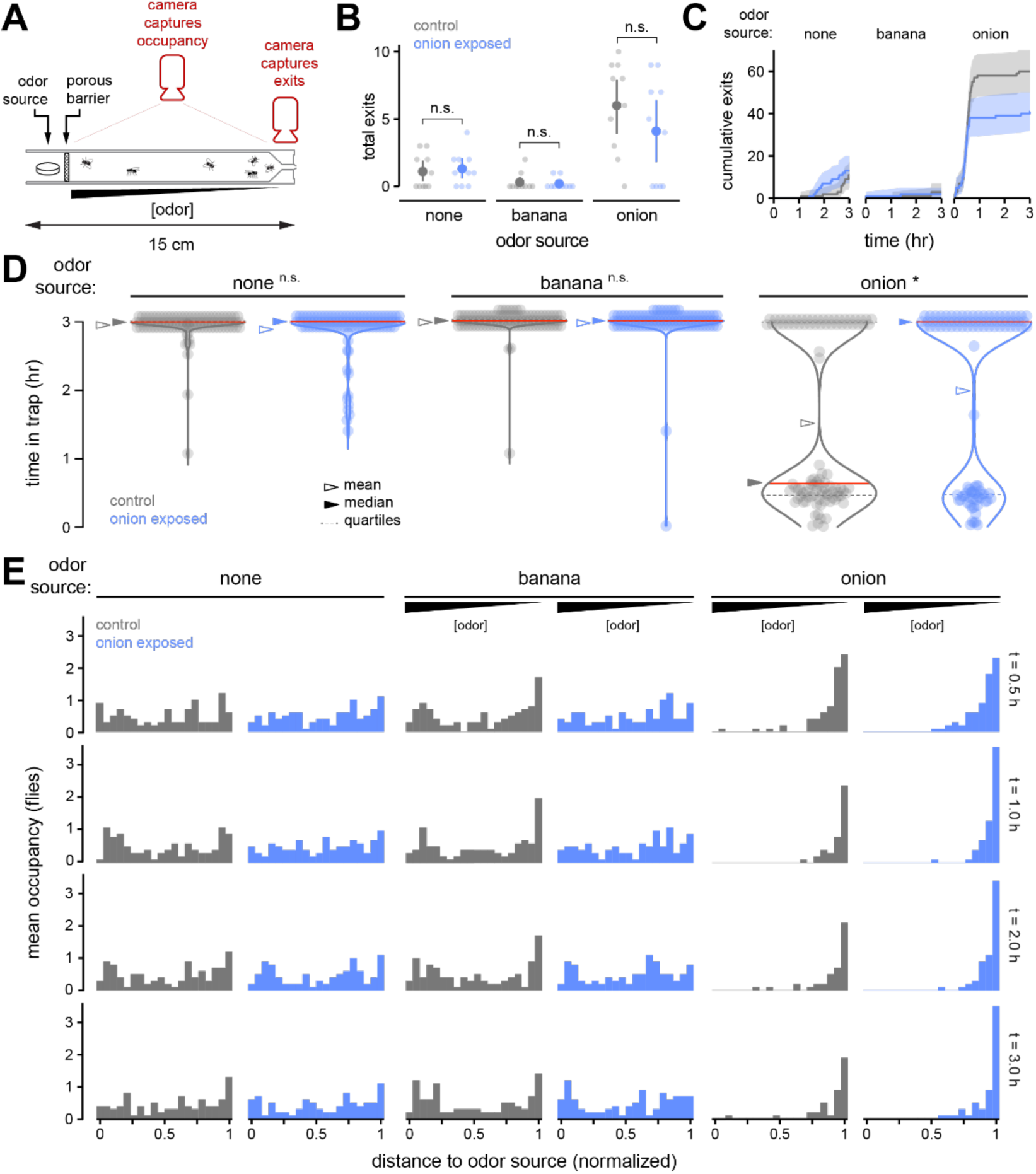
Persistent exposure to onion odor in early life reduces behavioral avoidance of onion odor selectively. **(A)** Cross-sectional view of the exit assay. Reproduced from Figure 1D. The positions and exits of ten unstarved flies from the odorized environment were recorded in each experiment. **(B)** Total exits (mean and 95% CI, solid symbols) over three hours for control- (gray) or onion odor- (blue) exposed flies from environments odorized with either banana, onion, or no odor (control) (n = 10 experiments, light symbols). n.s., no significant difference (*p* ≥ 0.05), two-tailed permutation t-test with Bonferroni correction. **(C)** Cumulative exits from environments odorized with each natural source across all experiments (n = 10) for each condition in **(B)**. Error envelope is the 95% CI bootstrapped across experiments. **(D)** Distributions of time spent in each environment for all animals (100 flies) across all experiments (n = 10) in each condition. Flies that never exit the environment during the assay period are assigned a dwell time of three hours. Triangles mark the mean (open) or median (solid) dwell times in each condition. \**p* < 0.05, n.s., no significant difference (*p* ≥ 0.05), two-tailed permutation t-test with Bonferroni correction. **(E)** Mean occupancy along the normalized length of each trap across all experiments (n = 10) for each condition in **(B)**, at 0.5, 1, 2, and 3 hours after flies are introduced into each odorized environment.

Inspection of the spatial distribution of flies in each odor gradient over time showed that, irrespective of their prior experience, flies avoided higher concentrations of onion odor. In comparison, flies were more uniformly distributed in the environment when it was non-odorized or odorized with banana (Figure 5E). These observations suggest that exposure to onion odor in early life does not strongly affect the sensitivity or ability of flies to detect onion odor. As the experiment progressed, larger numbers of onion odor-exposed flies compared to control flies accumulated at the far end of the onion odorized environment, away from the odor source (Figure 5E); this difference reflects the reduced number of exits from the onion environment by onion odor-exposed flies compared to controls. We conclude that chronic exposure to an aversive odor in early life can modestly reduce its behavioral aversiveness but is not sufficient to convert it into an attractive stimulus. Thus, behavioral responses to odor are not completely flexible; they can be shifted by odor experience, but this plasticity is bounded by the naïve olfactory preferences of the animal.

### PN sensitivity to familiar or unfamiliar odors is unaffected by chronic experience with a natural odor in early life

Second-order projections neurons (PNs) are the sole source of olfactory input to higher-order brain areas that process odor information. Some prior studies have shown that chronic exposure to high concentrations of monomolecular odors in early life reduces PN activity selectively in response to the familiar odor^17,18^. To evaluate the impact of early life experience with natural odors on PN odor representations, we used two-photon functional calcium imaging to compare PN activity in flies chronically exposed to banana odor in early life with activity in control-exposed flies. As before (Figure S1), we expressed the genetically encoded calcium indicator GCaMP6f in the majority of PNs (using the *NP0225-Gal4* driver) and confirmed that chronic exposure to banana odor in early life elicits behavioral plasticity in this specific genotype (Figure S3A-C). In each experiment, we imaged from PN axon terminals in a single imaging plane (∼17 um below the dorsal boundary of the calyx) through the mushroom body calyx and recorded PN responses to varying concentrations of each of three odors – banana, wine, and 2-butanone (Figure 6A, D). As controls for odor specificity and the responsiveness of the preparation, PN responses to an empty vial, to water, and to a mix of monomolecular odors were also measured in every experiment (Figure S3D).

**Figure 6:**
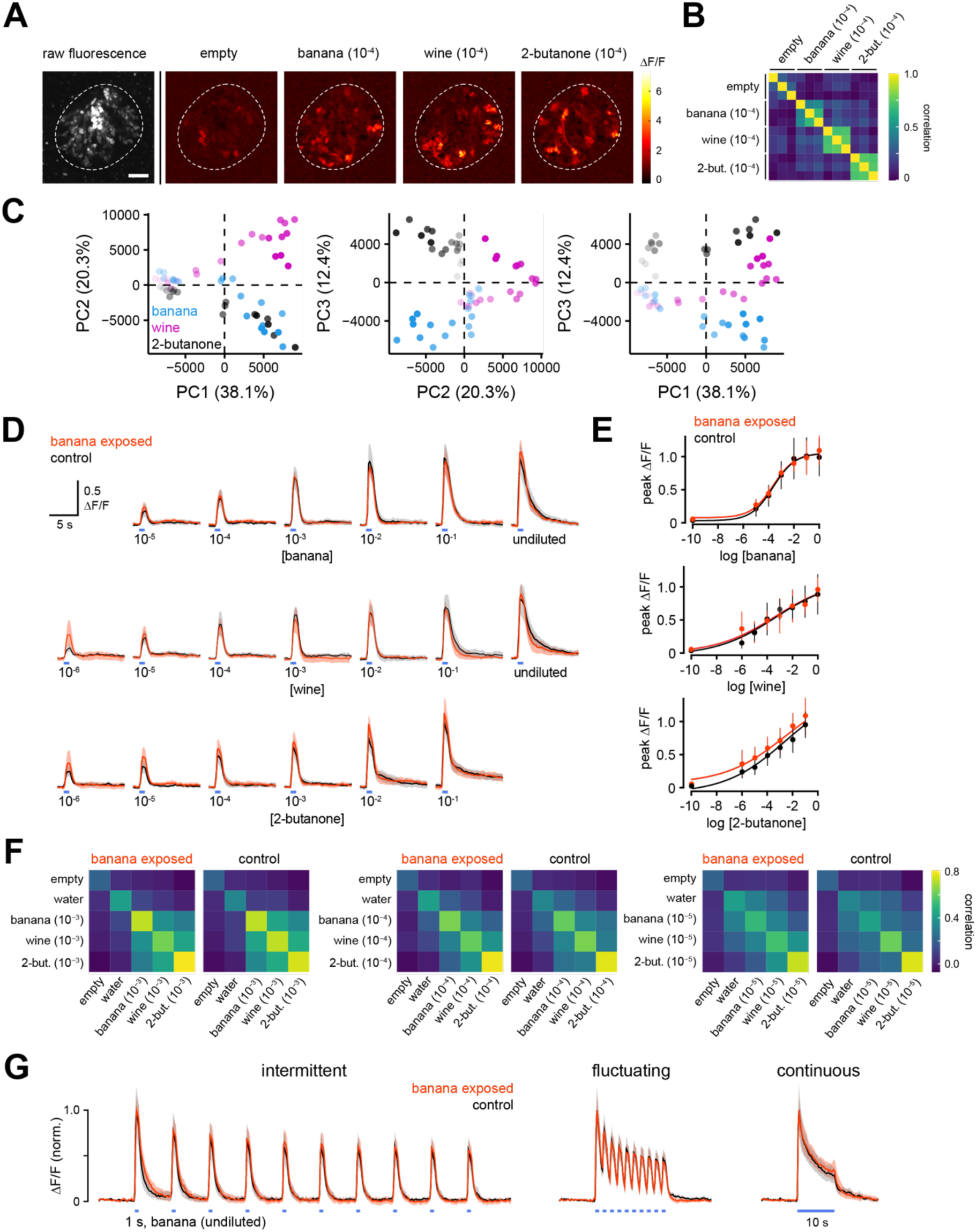
The sensitivity of PN responses to familiar or unfamiliar odors is unaltered by chronic experience with banana odor in early life. **(A)** Peak ΔF/F odor-evoked response patterns in PN axon terminals in an example imaging plane through the mushroom body calyx (white dashed line) of flies expressing GCaMP6f in a large subset of PNs. Grayscale image (left) shows raw fluorescence. Images (right) are trial-averaged peak responses (three trials/stimulus) to the odors of empty vial, banana, wine, or 2-butanone, each diluted 10^-4^ in water. Scale bar, 10 µm. The fly genotype was *yw/+; NP225-Gal4/*+; *20x-UAS-IVS-Syn21-OpGCaMP6f-p10*/+; +. **(B)** Pairwise Pearson correlation across pixels of peak PN response patterns evoked by different odor stimuli in **(A).** Each small block is the similarity between the response patterns elicited in individual trials of the indicated stimulus. **(C)** Principal component analysis of odor-evoked response patterns in PN terminals across all stimuli in the concentration series for banana, wine, and 2-butanone (same fly as in **(A), (B)**). Odor stimuli are as in **(D)**. Numbers in parentheses indicate the percentage of the variance in the data accounted for by each principal component. **(D)** Change in fluorescence over time (mean and 95% CI) in PN axon terminals in response to increasing concentrations of banana odor, wine odor, or 2-butanone odor in banana odor- (red) and control- (gray) exposed flies (n = 5–11 flies, banana odor-exposed; n = 5—9 flies, control-exposed). Blue bar indicates time of 1-s odor pulse. **(E)** Dose-response curves plotting peak PN response amplitudes from **(D)** (mean and 95% CI) at each concentration of each odor. **(F)** Mean pairwise Pearson correlation of peak PN response patterns evoked by different odors, at three different stimulus intensities, in banana odor- and control-exposed flies. Odor relationships, as measured by correlation in PN activity, are not significantly different between banana odor- and control-exposed flies for any pair of odors at any concentration (*p* ≥ 0.05, two-tailed permutation t-test with Bonferroni correction). **(G)** Change in fluorescence over time (mean and 95% CI) in PN axon terminals of banana odor- (red) and control- (gray) exposed flies (n = 7-8 flies) in response to banana odor presented with three different temporal structures (stimulus timing indicated by blue bars). To allow direct comparisons of response dynamics, bulk ΔF/F signals were normalized to the peak amplitude in every trial. Odor stimuli comprised of 10 s of undiluted banana odor presented as a 0.1 Hz train at 10% duty cycle (intermittent); a 0.5 Hz train at 50% duty cycle (fluctuating); or a sustained 10-s pulse (continuous).

The spatial patterns of peak odor-evoked responses across PN boutons were reliable and odor-specific: patterns of PN bouton activity from trials of the same odor were highly correlated to one another compared with PN activity from trials presenting different odors (Figure 6B). Principal component analysis on the spatial patterns of PN activity elicited across all stimuli in all three odor concentration series (in an individual fly) showed that, whereas the first principal component (PC) captured mostly variation from stimulus intensity, as expected, PN response patterns elicited by the three odors were well separated by their weightings on PC2 and PC3 (Figure 6C). The separability of PN representations of different odors increased with stimulus intensity (Figure 6C, S3E). Thus, odor-evoked activity in PN boutons reliably encodes stimulus identity.

The mean peak PN response amplitude, computed in an ROI circumscribing all PN boutons in the imaging plane for each fly, increased with concentration for all odors. Comparing PN response amplitudes in banana odor-exposed flies and control flies, we observed no significant differences in PN sensitivity to any odors, including to the familiar odor banana (Figure 6D-E). Chronic exposure to banana odor also did not significantly affect the relationship between any odors, including banana odor, in the representational space defined by PN bouton responses (Figure 6F). Odor relationships were determined by computing the mean pairwise correlation between peak PN activity patterns, evoked by each odor at approximately matched intermediate intensities, in banana odor-exposed and control flies.

Finally, we asked whether the dynamics of PN activity in response to odor stimuli of varying temporal structure are affected by chronic odor exposure in early life. Calcium signals in PN axon terminals were recorded in banana odor-exposed and control flies in response to a combined 10 seconds of banana odor presented with the following temporal structures: a 0.1 Hz train at 10% duty cycle (intermittent), a 0.5 Hz train at 50% duty cycle (fluctuating; Figure S3F), or a sustained 10-s pulse of odor (continuous) (Figure 6G). These stimuli probe how PN responses adapt when encountering odor stimuli that vary on different timescales. Bulk PN bouton calcium signals were normalized to the peak amplitude elicited in each trial to visualize the relative adaptation of the PN response over the course of the stimulus. We observed that PN boutons responded with distinct and reliable dynamics to each of the three stimulus structures, but the dynamics of PN output response amplitudes were indistinguishable between banana-exposed and control flies (Figure 6G). Taken together, we conclude that prior experience with natural odors does not strongly affect the sensitivity of olfactory PNs, to either familiar or unfamiliar odors, nor does it change the dynamics of PN output responses. These results indicate that the earlier and more frequent entry by banana odor-exposed flies into the banana trap are not accounted for by changes in either the sensitivity or overall level of activation driven by the familiar natural odor at early stages of sensory processing.

### Odor experience-dependent behavioral plasticity requires the presence of food during odor exposure

Given that olfactory experience that reliably alters behavioral attraction to odor does not strongly affect PN sensitivity, behavioral plasticity triggered by exposure to natural odors likely arises from changes in the olfactory circuit downstream of PN output. Experience-dependent behavioral plasticity elicited by chronic manipulation of the sensory environment has typically been interpreted as a form of non-associative learning that requires only passive long-term exposure to specific stimuli, without reinforcement or feedback^1,2,5,17,18^. Another possibility, however, is that continual experience with a specific natural odor while inhabiting an environment that robustly supports survival and reproduction could lead to formation of an association between the odor and the environment, increasing the attractiveness of the odor to the fly.

To disambiguate between these possibilities, we constructed a custom device for rearing flies that allowed us to temporally disassociate chronic exposure to a specific odor from the presence of food, since accessible, nutrient-rich food is a key feature of a high-quality environment. Groups of ∼500 flies were reared for two days in an acrylic cylinder (∼280 cm^3^) that could be rapidly odorized and de-odorized (Figure 7A). A platform fitted tightly against the base of the cylinder moved slowly (∼0.5 mm/s) on a linear track between two positions ∼7 cm apart – one in which the open base of the cylinder snugly abutted a smooth plastic surface, and the other in which it was tightly opposed to the surface of a well of standard molasses-cornmeal fly food (Figure 7A). Flies reared in this device were exposed to alternating 30-min epochs of banana-odorized and clean (unodorized) air (Figure 7B). Photoionization measurements of the odor environment in the cylinder confirmed that 30-min odor bouts were stable in amplitude and dynamics across 24 hours (the odor source was refreshed daily) (Figure 7C). In the paired condition, flies were positioned above the food during the epochs when banana-odorized air flowed through the cylinder and above the plastic surface during the epochs of non-odorized air, and vice versa for flies in the unpaired condition. In this way, both experimental groups of flies (paired and unpaired) were exposed to banana odor and given access to food for the same amount of time in total, with the only difference being whether food was made available to each group during the banana-odor epochs (paired) or the air epochs (unpaired) (Figure 7B). For both the paired and unpaired condition, a separate group of control flies were also reared in parallel using the same procedures, except clean unodorized air was substituted for banana-odorized air.

**Figure 7:**
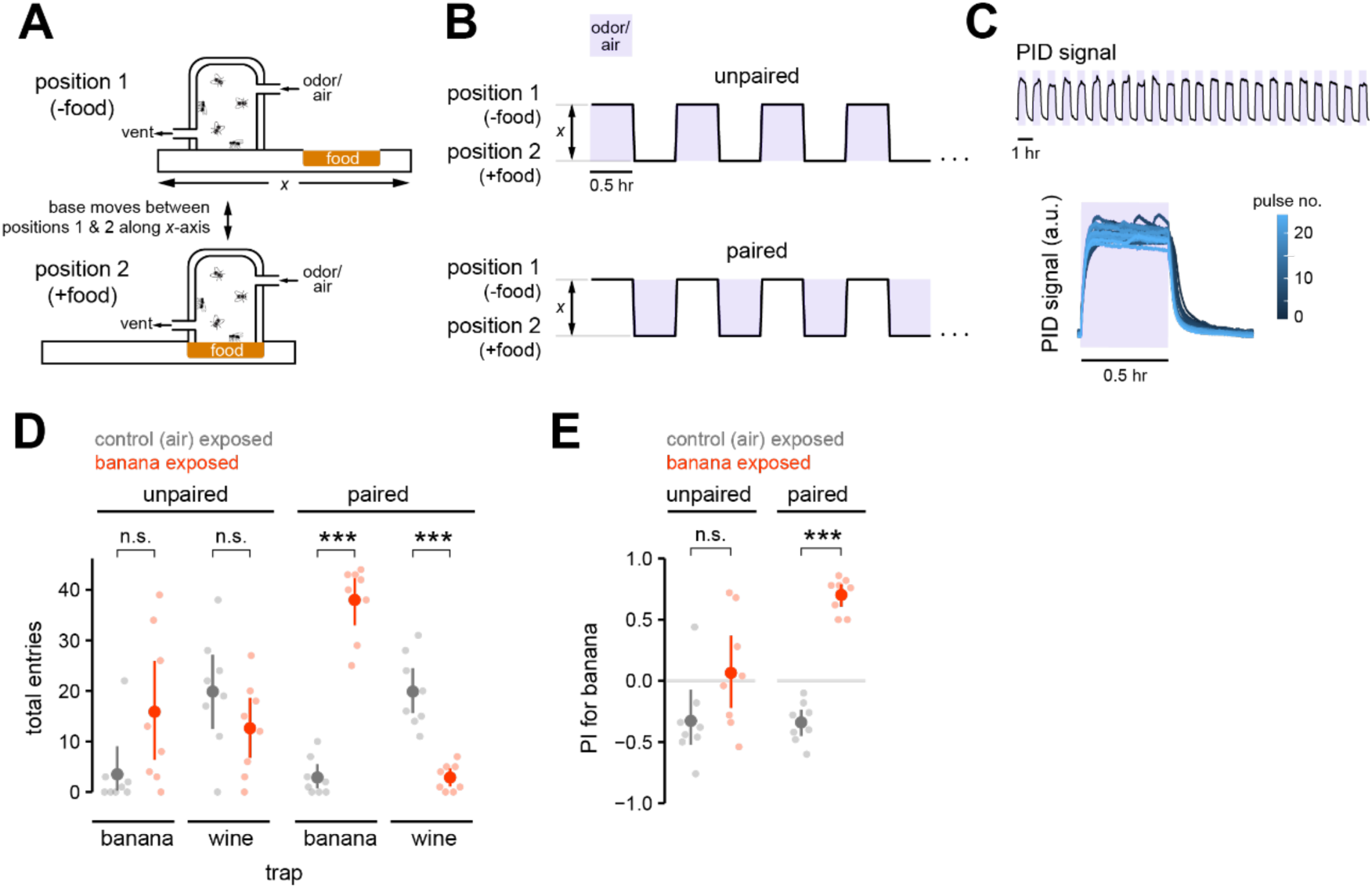
Increased behavioral attraction evoked by chronic exposure to natural odors requires the presence of food during odor exposure. **(A)** Schematic of custom device for rearing flies that allows temporal disassociation of long-lasting odor exposure and access to food. The base of the device, which sits tightly apposed to the cylinder housing the flies, shifts between two positions: one which provides a smooth, plastic base (position 1, -food) for the cylinder and the other which provides access to a well of food (position 2, +food). Humidified air, which is odorized if desired, flows constantly through the housing cylinder. **(B)** Unpaired (top) and paired (bottom) odor exposure. In the paired exposure condition, banana odor flows through the housing cylinder containing the flies only during the 30-min epochs it is apposed to food (position 2, +food). In the unpaired exposure condition, banana odor flows through the housing cylinder only during the 30-min epochs it is apposed to a plastic base (position 1, -food). Flies are reared in the custom device for 48 hrs. Flies in either exposure condition (paired or unpaired) experience banana odor and have access to food for equal amounts of time. Control-exposed flies are treated equivalently except that clean air, rather than banana odor, is used. **(C)** Photoionization measurement of the concentration of banana odor in the housing cylinder over 24 hours (top). The odor pulses for the 30-min odorized portion of every hour are overlayed (bottom panel), aligned to the onset of the TTL trigger for the odor valve, and color-coded by time. The natural odor source is refreshed after 24 hours. **(D)** Total entries (mean and 95% CI, solid symbols) over 24 hours into the banana- or wine-odorized trap by control- (gray) and banana odor- (red) exposed flies (n = 8 experiments, light symbols), exposed using the unpaired or paired procedure. \*\*\**p* < 0.001, n.s., no significant difference (*p* ≥ 0.05), two-tailed permutation t-test with Bonferroni correction. **(E)** Preference index for banana odor (mean and 95% CI, solid symbols) for each odor exposure condition in **(D)**. \*\*\**p* < 0.001, n.s., no significant difference (*p* ≥ 0.05), two-tailed permutation t-test with Bonferroni correction.

Flies were reared for two days in the device, starved for one day, and then tested for their preferences for wine or banana odor using the free-flight trap assay. Flies reared in our custom device tended to enter the traps with increased latency, so we extended the length of the assay period and evaluated the number of entries occurring within 24 hours. As expected, air-exposed control flies, from either the paired or unpaired condition, exhibited the same odor preference as naïve flies: they reliably preferred the odor of wine to that of banana (Figure 7D, Figure 4D-F), demonstrated by a greater number of entries into the wine trap compared to the banana trap. However, banana odor-exposed flies from the paired condition, which experience banana odor in the presence of food, strongly preferred banana odor over wine odor (Figure 7D-E). This result shows that paired odor exposure in the custom rearing device increased behavioral attraction to the odor, recapitulating the behavioral plasticity observed under more naturalistic conditions where the odor source is simply placed into the growth environment of the fly (Figure 4D-F). However, banana odor-exposed flies from the unpaired condition, which chronically experience banana odor without the presence of food, did not exhibit an increased attraction to banana odor and were similarly attracted to the odor of banana and wine (Figure 7D-E).

Together, these data demonstrate that passive odor experience alone is insufficient to elicit changes in olfactory behavior towards the odor. The change in olfactory preference elicited by chronic exposure to banana odor depends on additional features of the environment beyond just the prevalence of banana odor; the presence of food during odor exposure is also required. These results strongly suggest that behavioral plasticity elicited by chronic exposure to natural odors in early life represents a form of associative learning; its relationship to highly studied forms of classical conditioning in *Drosophila* is an important open question.

## DISCUSSION

### The role of odor experience in shaping the fly olfactory system in naturalistic conditions

The goal of this study was to better understand the role of odor experience in shaping the olfactory system of flies living in natural environments. Past work investigating the impact of odor experience on fly olfaction chronically exposed flies in early life to intense monomolecular odorants, olfactory stimuli that are universally aversive to naïve flies and are very rarely found in natural environments. These experiments concluded that the primary effect of chronic odor exposure was to reduce behavioral responses towards familiar odors^13,17,18,23,24^. Such behavioral habituation was determined to be the result of stimulus-selective changes in olfactory gain in PNs, implemented by neuron-specific increases in GABAergic inhibition^17,18^, and acted to match the sensitivity of the fly’s olfactory system to the local distribution of odors in the environment^16,37^.

We asked whether this framework describes how flies are affected by constant and long-lasting exposure in early life to natural odor sources that may be overrepresented in their local environment, including odors that are innately attractive to naïve flies. If behavioral plasticity elicited by chronic odor exposure were principally driven by long-term sensory adaptation in PNs, we expected to observe decreased aversion to innately aversive natural odors and decreased attraction to innately attractive natural odors. Instead, we found that chronic exposure to a natural odor in early life results in increased behavioral attraction selectively to the odor, irrespective of whether the odor is innately attractive or aversive to naïve flies. Thus, behavioral plasticity elicited by chronic experience with natural odors acts to promote acceptance and attraction to familiar odors experienced in early life.

### PN sensitivity is unchanged by long-lasting, persistent activation with natural odors

We found that odor-evoked output responses in PN axon terminals are not strongly altered by chronic exposure in early life to natural odors that strongly excite the fly olfactory system^38^ (Figure 6, Figure S1). In flies exposed to banana odor, PN response sensitivity was unchanged in response to both familiar and unfamiliar odor stimuli across a concentration range spanning several orders of magnitude (Figure 6D). PN response dynamics were also stable (Figure 6E). These results diverge from prior findings that chronic exposure of flies to high concentrations of the monomolecular odors carbon dioxide or ethyl butyrate reduces PN sensitivity selectively to the familiar odor, giving rise to behavioral habituation^17,18^.

Some of the differences between the current study and past studies on the impact of chronic odor experience on PN sensitivity may reflect methodological issues, for instance, the use of a different calcium indicator^39^ in functional imaging measurements. However, the differences may also reflect that chronic experience with different types of odor stimuli has varying long-term effects on olfactory physiology^38,40,41^. In past studies, animals were chronically exposed to monomolecular odors at estimated concentrations of ∼10^3^ to ∼10^5^ ppm in air^17,18,23,24,42–46^, whereas the headspace concentrations of the most abundant monomolecular volatiles common in natural odor sources like fruit typically range from ∼1 ppb to ∼10 ppm in air^47–49^. Even when all individual volatiles for a natural mixture are summed, the estimated upper bound of the total concentration of headspace volatiles for the vast majority of natural odor sources is ∼100 ppm in air^28,29,49^. For ripe banana^28,50^ and pineapple^29^, the estimated concentration of total headspace volatiles is ∼1-2.5 ppm or ∼250 bbp in air, respectively (using a vapor pressure estimate of ∼15.3 mm Hg at 25°C for all volatiles). Thus, an important difference in our study is that animals are exposed to significantly lower overall concentrations of odor volatiles. We note that recent studies have reported limited PN plasticity, or sometimes even slightly increased PN sensitivity, in flies chronically exposed in early life to monomolecular odors at low concentrations. These concentrations are within the range encountered in natural sources, but still strongly activate individual high affinity odorant receptors^40,51^.

Importantly, natural odor mixtures, such as wine or banana odor, can evoke overall levels of PN activity similar in strength to that evoked by an intense monomolecular stimulus (for example, 1% 2-butanone is ∼10^3^ ppm in air) (Figure 1G, and data not shown). This result indicates that differences in the overall extent to which different odor stimuli activate the olfactory system do not account for differences in their ability to trigger PN plasticity. Another key factor is that natural odors are usually complex blends of dozens of distinct monomolecular odorants^25,26,48,49^. Thus, the distribution of input activity to the olfactory system (e.g., how narrowly or broadly an odor stimulus acts across olfactory receptor neuron classes) elicited by monomolecular odors or natural odors may be an important factor contributing to how chronic olfactory excitation affects PN odor coding. However, our study demonstrates that olfactory behavioral plasticity elicited by persistent odor experience need not necessarily arise from changes in PN sensitivity.

Understanding the effects on olfactory function of chronic exposure to either monomolecular odors or to natural odors is important. The impact of persistent experience with specific natural odors provides insight into the adaptive functions of olfactory plasticity in the natural context in which olfactory systems evolved. However, human activity creates local environments where people and other organisms can be chronically exposed to artificially high concentrations of a particular monomolecular volatile. For instance, the National Institute for Occupational Safety and Health establishes guidelines for safe indoor exposure limits to specific volatiles from the perspective of acute human toxicity, in the range of ∼10^2^-10^3^ ppm in air for several hours for most volatiles^52^. However, the long-term consequence of unabated exposure to sub-lethal, but still high concentrations of monomolecular volatiles for olfactory physiology and behavior requires more investigation.

### The role of innate preference in odor experience-dependent behavioral plasticity

Generally, we found that odor experience in early life shifts olfactory preference towards increased behavioral attraction to the familiar odor. However, odor experience cannot shift olfactory preference sufficiently to convert behavioral aversion towards a naively aversive odor, such as onion odor, into behavioral attraction: onion odor-exposed flies, like naïve flies, never select the onion trap and still strongly prefer banana odor (Figure 2C). Also, in the exit assay, although onion odor-exposed flies are more willing to dwell in the onion-odorized trap (Figure 5D), they still avoid onion odor, preferring to occupy the side of the trap with the lowest concentration of odor (Figure 5E). On the other hand, rank odor preference is not immutable: for two innately attractive odors, wine and banana, the preference of naïve flies for wine odor over banana odor can be reversed by early life experience with banana odor (Figure 4). Our data suggest the existence of inherent boundaries to the degree that odor preference can be altered by naturalistic experience, and these boundaries are likely odor-specific. Odors emanating from a toxic or dangerous source are often innately aversive, and a hard limit to the degree that olfactory aversion can be modified by experience may preclude catastrophic outcomes for the animal. However, a switch in preference between two attractive odors signaling food may be adaptive, promoting the selection of the one associated with reliable food and/or reduced risk during foraging. The form of behavioral plasticity triggered by chronic exposure to a specific odor stimulus may depend on the innate meaning or valence of the stimulus to the fly, on the identity and degree of activation of specific olfactory receptor neurons by the stimulus, or on both.

### Chronic exposure to natural odors alters distinct aspects of odor-driven behavior

The free-flight odor trap assay models a simple foraging task, in which hungry flies search for food in the environment using odor as a cue. Trap entry is preceded by several stages of foraging behavior: first, flies locate from a distance and land on the surface of the trap; second, flies repeatedly approach and locally investigate the entrance apertures of the trap; and, third, flies make the final decision to walk through the entrance apertures and access the trap. By extracting the positions of individual flies on the trap over time from video data, we gained new insights into how different phases of this foraging behavior are affected by chronic experience with natural odors (Figure 3).

We observed that the earliest stages of odor search are unaffected by prior olfactory experience, with all groups of flies locating, landing, and occupying the platform of the banana-odorized trap for a similar amount of time in the first few minutes of the experiment. However, flies having long-lasting experience with banana odor in early life exhibited altered behavior at later stages of odor search: they investigated the entrance apertures of the trap, where odor concentration is highest, with a greater intensity, making earlier and more frequent approaches to the trap entrances. Flies repeatedly approached and sampled the entrance holes (Supplementary Video S1), often extending half their body length into the entrance aperture before withdrawing, briefly leaving, and returning to repeat the process. This series of behaviors suggests that passage through an entrance aperture into the trap carries a certain degree of risk. This risk may be overcome by repeated exploration of the trap entrances, increasing pressure in starved animals to locate energy resources, and/or prior knowledge about the odor emanating from the apertures.

The temporal profile of behavior is consistent with functional imaging measurements indicating that odor-evoked output from PNs is unaffected by chronic odor exposure. Irrespective of earlier olfactory experience, all groups of flies appear to detect and locate banana odor with comparable latencies, inconsistent with a significant impact of chronic odor exposure on olfactory behavioral sensitivity or habituation. Rather, odor experience elicits long-lasting changes in the flies’ olfactory exploratory behavior and alters the behavioral threshold for odor-mediated passage through the narrow trap entrances. These changes in foraging behavior suggest that long-lasting odor exposure could be teaching flies something about either the value and/or risk that they associate with the odor.

### Natural odor experience changes olfactory behavior through an associative process

The development of a novel rearing environment that allowed the temporal decorrelation of food access from odor exposure enabled us to evaluate whether food-related cues in the environment are important for behavioral plasticity triggered by natural odors^41^. Investigating the effects of long-term exposure to biased sensory environments on animal brains usually requires the provision of food in the environment, and past work investigating the effects of chronic exposure to monomolecular odors have necessarily been performed in the presence of food^17,18,23,24,40,43,45^. We discovered that the induction of behavioral plasticity by chronic odor exposure depends critically on the presence of food during intervals of odor exposure. Our results are consistent with a scenario where the temporal coincidence of odor and food over long timescales teaches flies that the familiar odor reliably signals the local availability of a nutrient-rich energy source, through an associative process.

Learning elicited by chronic odor experience may be related to, but is distinct from, well-established forms of associative learning like classical olfactory conditioning, a form of learning that has been powerfully dissected at the genetic and neural circuit level in *Drosophila*. Chronic olfactory experience-dependent plasticity differs in several ways, including: 1) flies are not food-deprived prior to the learning phase, whereas in classical appetitive odor conditioning, flies are strongly starved before training with sugar reinforcement; and 2) the pairing of odor and reinforcement (presumptively the food environment) occurs over long timescales (hours), whereas in classical appetitive odor conditioning, odor and sugar reinforcement are temporally paired within a seconds-to minutes-long window^53,54^. Both types of learning induce strong, long-lasting plasticity that persists for at least a day.

Thus, olfactory experience-dependent plasticity and classical olfactory conditioning are two lab-based paradigms that model odor-guided associative learning and prediction occurring on different natural time scales; they likely come into play in different behavioral contexts. For instance, whereas olfactory experience-dependent plasticity may be important for learning the reward landscape of an environment through extended exploration and foraging, classical associative learning may mediate fast or single-trial learning important for avoiding danger or obtaining transient rewards. A fascinating topic of future investigation is the degree to which olfactory experience-dependent plasticity, elicited under slower, naturalistic behavioral timescales, is mediated by mechanisms common or divergent from the well-studied circuitry mediating classical olfactory conditioning in the mushroom body.

Olfactory experience-dependent plasticity may also be viewed through the lens of perceptual learning, defined as a stimulus-selective enhancement of the ability of the flies to smell the familiar odor. On balance, our results argue against chronic experience with natural odors eliciting a change in sensitivity towards the odor: olfactory experience does not impact PN sensitivity to the familiar odor nor does it significantly alter its representational distance from other odors (Figure 6), and the behavioral latency to initially locate the odor is unaltered by experience.

However, chronic odor experience may affect the ability of the fly to recognize or otherwise interpret a familiar odor. An unavoidable feature of using natural odor sources is that the odor emitted cannot be exactly identical between exposure and testing. The pieces of fruit are slowly spoiling and/or fermenting over the days of odor exposure, and different pieces of fruit are used in exposure and testing. Furthermore, during odor exposure, additional odors are necessarily present in the growth environment, including the odor of fly food and odors from other flies at close range, which are not present during testing. This aspect of our experimental design was intentional and meant to emulate the types of olfactory conditions and tasks flies likely encounter in the natural world. An interesting possibility is that odor experience improves the ability of flies to generalize or group together variants of an ecologically important natural odor as an odor object that been previously encountered and is predictive of a reliable source of food. Indeed, in the auditory system, persistent exposure to single frequency tones paradoxically leads to poorer discrimination in the overrepresented frequency band, consistent with perception being more stable in the face of stimulus perturbations^19^. Such changes in perceptual judgment could also contribute to experience-dependent increases in exploration and entry into traps odorized with a familiar source.

The division between perceptual learning and associative learning ultimately may reflect historical distinctions, rather than a true mechanistic dichotomy. Perceptual learning most commonly occurs in the context of training to a specific task, and it is well understood that both attention and reward have important roles in most forms of perceptual learning. Additionally, reward processing and reinforcement impact how the brain forms perceptual judgments in sensory-based associative learning tasks^55^. In rodents, perceptual learning in the olfactory system occurs in the context of reinforcement-based conditioning tasks^56–59^. Thus, chronic experience with natural odors may elicit selective changes in the perception of the familiar odor, in the value or risk associated with the familiar odor, or in both.

### Learning about odor environments is beneficial

The non-classical form of learning elicited by chronic experience with natural odors in early life may not act to principally predict rewards on fast timescales, but rather may be important for allowing the animal to associate specific odors with high quality environments or with a general state of well-being. From an ethological perspective, it is logical that neural sensitivity to familiar odors at early stages of olfactory processing are unaffected by olfactory experience. If an odor is associated with a positive environment, maintaining sensitivity to that odor would be beneficial by supporting its reliable detection, whereas sensory habituation would have an opposite outcome. Moreover, stability in odor representations at early stages of processing would allow long timescale experience-dependent learning to interact with short timescale classical associative learning, enabling the flexible reformatting of odor meaning in higher brain areas if local conditions shift rapidly. Our results extend the forms of learning we understand to occur in *Drosophila* to include non-classical associative learning at long timescales. Important future goals will be to understand the use and interaction of these different forms of learning during odor-guided adaptive behavior in natural contexts, as well as to elucidate the shared and divergent neural mechanisms underlying these different forms of olfactory learning.

## AUTHOR CONTRIBUTIONS

**Kristina V. Dylla:** Conceptualization, Methodology, Investigation, Validation, Formal analysis, Writing – Original Draft, Writing – Review & Editing, Visualization. **Thomas F. O’ Connell:** Methodology. **Elizabeth J. Hong:** Conceptualization, Methodology, Formal analysis, Writing – Original Draft, Writing – Review & Editing, Visualization, Supervision, Funding acquisition.

## ACKNOWLEDGEMENTS

We thank M. Dickinson for sharing *Canton-S* flies and members of the Dickinson lab for advice on fly tracking. We thank D. J. Anderson for sharing unpublished flies. We are grateful to D. A. Wagenaar and the Caltech Neurotechnology Core for collaborating on the design and construction of the odor trap cores and the custom fly rearing device. We thank V. Hauser and A. Ruiz Sandoval for assisting with fly food preparation, and D. Kim, B. G. Fang, J. Eden-Sung, and J. Saltzman for assistance with processing video data. We thank members of the Hong lab for their careful reading and comments on this manuscript. This work was funded by grants to E.J.H. from the NSF/CIHR/DFG/FRQ/UKRI-MRC Next Generation Networks for Neuroscience Program (NeuroNex Award #2014217) and the Shurl and Kay Curci Foundation. K.V.D. was supported by grants from the Della Martin Foundation and the German Research Foundation (DY 135/1-1). E. J. H. is a Chen Scholar of the Tianqiao and Chrissy Chen Institute for Neuroscience and a Clare Boothe Luce Professor of the Henry Luce Foundation.

## METHODS

### Resource Availability

#### Lead Contact

Further information and requests for resources should be directed to and will be fulfilled by the Lead Contact, Elizabeth J. Hong

#### Materials Availability

This study did not generate new biological reagents. Design schematics for the odor trap inserts and the custom rearing device are available upon request.

#### Data and Code Availability

- All functional imaging and behavioral videos will be made available to any researcher for the purposes of reproducing or advancing the results.
- Software in this study was adapted from existing code. All custom scripts have been deposited at GitHub and are publicly available as of the date of publication. The URLs are listed in the key resources table.
- Any additional information required to reanalyze the data reported in this paper is available from the lead contact upon request.

### Experimental Model and Subject Details

*Drosophila melanogaster* were raised on a 12:12 light:dark cycle at 25°C and 70% relative humidity. Flies were mostly raised on Nutri-Fly® Molasses Formulation (# 66-123, Genesee Scientific) supplemented with sucrose (1.6%, w/v) and propionic acid (0.45%, v/v). In some experiments, flies were raised on cornmeal/molasses food with composition: water (17.8 l), agar (136 g), cornmeal (1335.4 g), yeast (540 g), sucrose (320 g), molasses (1.64 l), CaCl_2_ (12.5 g), sodium tartrate (150 g), Tegosept (18.45 g), 95% ethanol (153.3 ml) and propionic acid (91.5 ml). These two food formulations have similar composition; in both cases, food was supplemented with dry yeast. Behavioral results were indistinguishable between the two food formulations. Behavioral experiments were performed in mixed populations of male and female flies of wild-type strain Canton-S. Functional imaging experiments were performed in female flies with genotype: *yw*/+*; NP225-Gal4*/+*; 20x-UAS-IVS-Syn21-OpGCaMP6f-p10*/+.

### Method Details

#### Fly stocks

The original stock of wild-type strain Canton-S was from M. Heisenberg lab and was a gift from M. Dickinson lab (Caltech). *NP225-Gal4* was acquired from the Kyoto DGGR Stock Center (DGRC: 112095). *20x-UAS-IVS-Syn21-OpGCaMP6f-p10* was a gift from D. Anderson lab (Caltech).

#### Odors

Odor sources used in this study were: apple cider vinegar, red wine, banana, pineapple, mango, onion, 2-butanone (Cat. #360473, Sigma-Aldrich), 2-pentanone (Cat. #46211, Sigma-Aldrich), E2-hexenal (Cat. #158131000, Sigma-Aldrich), and pentyl acetate (Cat. #109584, Sigma). Fresh banana (yellow ripe) and onion were purchased weekly from local grocery stores; for pineapple and mango, thawed frozen chunks were used. Apple cider vinegar and red wine were purchased from the grocery store. All fruits/vegetables were used without peel. For chronic odor exposure, the odor sources were narrow polystyrene *Drosophila* vials (25 x 95 mm) filled ∼1/3 (30 mm height) with small chunks of banana, onion, or pineapple, and sealed with mesh so that flies could not access the vial. For testing in the free flight trap-based assay, 2 ml of liquid odor sources or ∼1 cm^3^ solid odor sources were used. For testing in the exit assay, solid onion or banana pieces of ∼2 cm^3^ were used. For functional imaging studies, 2 ml of liquid odor sources or ∼2 cm^3^ solid odor sources were used. For the banana odor concentration series, undiluted banana was 2 ml of mashed banana, which allowed v/v dilutions. All odor dilutions were v/v in water (ddH2O) and were prepared fresh every day.

#### Chronic odor exposure

*Standard procedure.* Groups of ∼500 flies (< 24 hours post-eclosion) were transferred to fresh food bottles (∼237 ml round-bottomed, polypropylene) supplemented with dry yeast. A mesh- sealed vial containing the odor source was inserted into the bottle. The control-exposed group was treated the same way but was housed with an empty vial. Flies were maintained for two days in ambient room conditions, a few meters apart from each other to avoid odor cross-contamination. Odor sources were not refreshed during this period.

*Paired and unpaired exposure.* A custom device for rearing flies that allows temporal disassociation of long-lasting odor exposure and access to food was designed and constructed in collaboration with Daniel Wagenaar (Caltech Neurotechnology Laboratory). An acrylic cylinder (6 x 10 cm, ∼280 cm^3^ volume) with an open base, which housed the flies, was positioned tightly against an acrylic platform (29.5 x 7.5 cm), which could be moved bidirectionally on a set of rails along its long axis (*x*-axis in Figure 7A) using a bipolar stepper motor (NEMA 17HS4401; driver: A4988, HiLetgo). The acrylic platform contained two circular cutouts (diameter 6 cm), in each of which rested a plastic Petri dish (Falcon #351007, 6x 1.5 cm). The Petri dishes were completely filled with fly food and the surface of the food was leveled, such that all parts of the platform surface fit tightly against the base of the cylinder. A tight fit between the cylinder and the entire length of the platform was important to prevent loss of flies from the cylinder. Access to food was controlled by moving the platform to position one of the Petri dishes directly under the housing cylinder (+ food; position 2 in Figure 7A). We alternated between making each of the two Petri dishes available to flies in order to reduce the amount of time the food was exposed to air; excessive exposure to air resulted in dehydration and shrinkage of the food. In the “— food” condition, the solid plastic of the platform was positioned under the cylinder (position 1). Positions 1 and 2 were ∼7.5 cm apart. The platform base was moved slowly (∼0.5 mm/s) to prevent injury to the flies; transitioning from position 1 to position 2 (or vice versa) took ∼2.5 min. The full device comprised two acrylic cylinders for housing flies and two platforms holding the food, situated side-by-side. Both platforms were moved by the same stepper motor to allow rearing of odor-exposed and control-exposed flies in parallel.

The housing cylinder could be rapidly odorized and de-odorized. A humidified carrier stream (200 ml/min; 60-70% relative humidity) and an odor stream (100 ml/min) were combined and introduced into the cylinder, for a total constant airflow of 300 ml/min that entered the cylinder through a port (∼5 mm diameter) near the top and exited through a vent located near the bottom of the cylinder on the opposite side. For the banana odor exposure group, the odor stream switched between an empty 20-ml vial (for non-odorized epochs) or one containing a ∼2 cm^3^ piece of banana (for odorized epochs). For the control odor exposure group, the odor stream switched between two empty 20-ml vials.

Groups of ∼500 flies (< 24 hours post-eclosion) of mixed sex were exposed to banana odor (or clean air in the control group) using either paired or unpaired procedures (see Results; Figure 7B) for 48 hours. After 24 hours, the odor source was replaced with a fresh sample to ensure stable odor levels, and the fly food was also replaced with a fresh plate. Paired and unpaired odor exposure experiments were interleaved whenever possible. Odor concentrations inside the rearing cylinder were measured over 24 hours with a photoionization detector (200B miniPID, Aurora Scientific), sampling at 1 Hz. Coordination of the motor driving platform movement with the valves regulating odor delivery was controlled using an Arduino UNO (Arduino.cc) microcontroller board. Odor valves were switched to the next state when movement from one position to another initiated.

#### Starvation

Flies were deprived of food prior to testing in the free flight trap-based assay and prior to functional imaging experiments. Odor- or control-exposed flies were transferred to empty food bottles containing only moistened lab tissue. Flies were transferred from the exposure bottle to the starvation bottle without anesthesia and were also counted during this process. Wildtype flies were starved for 24 hr. Since *yw*/+; *NP225-Gal4*/+; *20x-UAS-IVS-Syn21-OpGCaMP6f-p10*/+ tend to survive longer without food compared to wildtype (Supplementary Figure S3A), we increased the starvation time for *yw*/+; *NP225-Gal4*/+; *20x-UAS-IVS-Syn21-OpGCaMP6f-p10*/+ flies in behavioral experiments to 30.5 hr (Supplementary Figure S3B-C). For functional imaging, *yw/+; NP225-Gal4*/+; *20x-UAS-IVS-Syn21-OpGCaMP6f-p10/*+ flies were starved between 24 and 30.5 hr.

#### Behavioral assays

*Free flight trap-based assay.* Groups of 50 starved flies of mixed sex were released into insect cages (45 x 45 x 45 cm; # 4S4545, BugDorm) containing two odor traps, positioned ∼18 cm apart on the floor of the cage. The exceptions were that, when testing flies exposed to pineapple or when testing *NP225>GCaMP6f* flies, groups of 70 flies were tested, because overall entry rate was lower in these conditions. Each trap was constructed from a 20-ml borosilicate glass scintillation vial, fitted with a white polyethylene cap (# 333714, Fisher Scientific). A ring (outer diameter, 95 mm; inner diameter, 24 mm) constructed from sturdy white foamboard was tightly fitted around the vial cap (diameter, 24 mm) such that the surface of the cap was level and continuous with the surface of the ring, creating a ∼10-cm circular surface on which flies could land and explore the region surrounding the trap entrances. Five circular holes (1.65 mm diameter), spaced ∼4 mm apart, were created in the center of the polyethylene cap by pushing a 16-gauge needle through the cap. A cylindrical insert (15 mm dia. x 50 mm height) fabricated from transparent acrylic was centered inside the vial. The outer diameter of this trap core (15 mm) was matched to the inner diameter of the mouth of the scintillation vial, so that the insert fit snugly in the vial. A funnel-shaped chamber (15 mm dia. mouth x 7 mm height) was bored out of the top of the cylindrical insert, and it narrowed into a stem with inner diameter ∼2 mm and length ∼4 mm. A trimmed pipette tip was fitted on the tip of the stem, further narrowing the funnel to ∼1.5 mm. Flies that passed through the funnel entered a cylindrical chamber (9.5 mm dia. x ∼40 mm height) containing the odor source. The narrowing of the funnel exit discouraged flies from re-entering the funnel stem. The stem was imaged from the side to capture entries into the trap. Flies were usually allowed to investigate the traps in free flight inside the cage for three hours, during which their positions on the trap platforms and entries into the traps were recorded. For the paired and unpaired odor exposure experiments in Figure 7, total trap entries were counted 24 hours after releasing flies because of the delayed onset of trap entries for flies grown in the custom rearing device. The reasons for delayed entry into the traps are not known. The assignment of odor sources (or water) to each of the two trap positions in each cage, and the assignment of exposure groups to different testing cages, were pseudorandomized across experiments.

*Exit assay.* Groups of ten unstarved flies of mixed sex were introduced into a clear cylindrical tube (15 dia. x ∼140 mm). A cotton plug on one end prevented flies from existing the chamber. The other end was fitted with the same custom fabricated trap core as described for the regular odor traps, creating a funnel-shaped exit through which flies could exit the chamber (Figure 5A). Newly introduced flies were first allowed to acclimate and randomly distribute in the cylindrical chamber for ten minutes. Then, the odor source (onion, banana, or none) was placed close, but not touching, the other side of the cotton barrier. The positions of flies in the cylindrical chamber, as well as individual exits from the chamber, were recorded for three hours. The assignment of odor sources to testing chambers at different positions in each cage and the assignment of exposure groups to different testing cages were pseudorandomized across experiments.

*Survival experiments.* Following 48 hours of chronic odor exposure, and then 24 hours of wet starvation, groups of 50 flies were briefly anesthetized with CO_2_ and transferred to petri dishes (10 x 1.5 cm) lined with Whatman paper. Video of flies was recorded from overhead for 33 hours, by which time all flies were dead.

*Video recordings.* Movies were captured using multiple webcams (HD Pro Webcam C920, Logitech) at each trap (Figure 1A), at a frame rate of 5 Hz and resolution of 640 x 480 pixels. A custom modification of the Multi tracker package (https://github.com/tom-f-oconnell/multi_tracker) was used for video acquisition.

#### Functional imaging

*Odor delivery.* Odors were delivered essentially as previously described^60^. A constant stream of charcoal-filtered air (2 L/min) was directed at the fly. When no odor was delivered, ten percent of the airstream (200 ml/min) was routed through the “normally open” port of a three-way solenoid valve (ASCO 411-L-1324-HVS, NISCO, Inc., Duluth, GA) and passed through the headspace of an empty 20-ml glass scintillation vial, before rejoining the main carrier stream (1800 ml/min). For odor delivery, an external trigger switched the valve to the ‘normally closed’ position and the 200 ml/min airstream was redirected through the headspace of a scintillation vial containing the odor source, before rejoining the carrier stream. The 200 ml/min control or odor streams were carried by tubing of matched lengths and rejoined the carrier stream at the same point along the carrier tube (I.D., ∼5 mm), approximately 10 cm from the point of exit. The end of the carrier tube faced the front of the fly and was positioned ∼2 cm away. Odors were vented by a vacuum line connected to a funnel that was positioned behind the fly. Flow rates were controlled using mass flow controllers (MC series, Alicat Scientific).

*Two-photon calcium imaging.* In vivo functional calcium imaging was performed essentially as previously described^61^. In brief, female flies were briefly anesthetized on ice (<15 s) and head-fixed with the head tilted ∼70° downwards and the posterior plate of the head positioned as level as possible, 90° with respect to the imaging axis. Cuticle, trachea, and fat were removed to expose the posterior surface of the brain and underlying mushroom body. The open brain was flooded with *Drosophila* saline (103 mM NaCl, 3 mM KCl, 5 mM N-Tris(hydroxymethyl)methyl-2-aminoethane-sulfonic acid, 8 mM trehalose, 10 mM glucose, 26 mM NaHCO3, 1 mM NaH2PO4, 1.5 mM CaCl2, and 4 mM MgCl2; pH 7.3, osmolarity adjusted to 270 – 275 mOsm), and the antennae and maxillary palps remained dry below the imaging chamber, accessible to the olfactometer carrier tube. The proboscis and legs were immobilized with UV glue, and the M16 muscle was severed, to reduce movement. Naïve PN odor responses were recorded in unstarved flies using regular saline. For measurements of PN odor responses in flies chronically exposed to odor, flies were starved for 24 hours after odor exposure, using the same procedure as for behavioral experiments. Calcium imaging was performed using sugar-free saline (regular saline in which glucose and trehalose were replaced with 18 mM ribose).

Two-photon GCaMP6f fluorescence was excited with 925 nm light from a Mai Tai DeepSee laser (Spectra-Physics, Santa Clara, CA). Images were acquired with a 20X water immersion objective (Olympus XLUMPLFLN20XW, 1.0 NA) on a two-photon microscope equipped with galvo-galvo scanners (Thorlabs Imaging Systems, Sterling, VA). Imaging was performed at a frame rate of 4.5 Hz at a resolution of 256 × 256 pixels (73.43 x 73.43 μm^2^). The collection filter was centered at 925 nm with a 50 nm bandwidth. The microscope setup was housed in a lightproof box. All experiments were conducted at room temperature (∼21°C). The brain was constantly perfused by gravity flow (∼2 - 3 ml/min) with saline oxygenated with carbogen (95% O_2_ / 5% CO_2_).

The imaging plane through the mushroom body calyx was chosen to sample a large number of projection neuron axon boutons while avoiding the primary axonal tracts of the PNs, typically ∼17-18 µm below the dorsal limit of the calyx. For measurement of naïve PN odor responses in Figure 1G, stimuli were presented in a different random order in each experiment. For measurement of PN odor responses in Figure 6, each stimulus block started with measuring the response to water, then odor stimuli were delivered with progressively increasing concentrations, alternating across different odors (banana, wine, 2-butanone) at each concentration step. Odor stimuli were delivered from sources diluted in the vial as follows: banana odor: 10^-5^, 10^-4^, 10^-3^, 10^-2^, 10^-1^, undiluted; red wine: 10^-6^, 10^-5^, 10^-4^, 10^-3^, 10^-2^, 10^-1^, undiluted; and of 2-butanone: 10^-6^, 10^-5^, 10^-4^, 10^-3^, 10^-2^, 10^-1^. The presentation order of odors (banana, wine, 2-butanone) was pseudorandom in each experiment. We attempted to measure responses to concentration series of all three odors sin every fly, but, in some flies, only two were recorded. Responses to an empty odor vial were recorded both prior to and after the odor panel to evaluate for odor contamination. The responses shown for the “empty” stimulus in the figures were from the first presentation of the empty odor vial. Responses to the monomolecular odor mixture (2-butanone, 2-pentanone, E2-hexanal, and pentyl acetate, each 10^-1^ in water) were measured at the end of each experiment to confirm the responsiveness of the preparation. In all experiments, odor stimuli were 1-s pulses presented in blocks of three trials, with a 40-s intertrial interval. The odor source for PN responses to banana odor of varying temporal structure in Figure 6G was a solid piece of banana (∼2 cm^3^). A single trial for each stimulus structure (intermittent, fluctuating, continuous) was collected per fly.

### Quantification and Statistical Analysis

#### Free-flight trap-based assay

*Trap entries*. Extraction of each event of a fly entering a trap from side-camera video data was performed with ImageJ (version 1.53f51; http://rsbweb.nih.gov/ij/). An ROI was defined in the center of the stem that connected the funnel of the trap insert and the core of the trap. Since flies were darker than the background of the trap, passage of a fly through the ROI resulted in a transient reduction in mean pixel intensity in the ROI. The mean pixel intensity in the ROI was computed for every frame of the movie. Temporal and intensity filters were applied in R (version 4.0.2; https://www.r-project.org/) in RStudio integrated development environment (version 1.4.1106; https://www.rstudio.com/), and the times of potential entries were extracted as the times of negative peaks in the plot of mean ROI intensity as a function of time. Filter settings were chosen with a bias towards false positives, and they were adjusted as needed for each movie. Visual inspection of the video data was performed to confirm each entry event. Manual auditing resulted in two types of corrections. Events were excluded when flies did not complete their entry into the trap core after passing through the stem or when flies passed through the ROI while walking on the exterior of the trap. Events were added when multiple flies that entered as a chain into the trap were counted as a single entry event. We very rarely observed cases where flies managed to escape from the trap core after they had entered; in such cases, the experiment was excluded from analysis. The total number of entries at the end of the three-hour experiment was computed for each experimental replicate, and the mean over the ten experiments for each condition was calculated. Cumulative entry curves were computed by summing entries at each time point across the combined data of the ten experimental replicates for each condition.

*Preference index.* The preference index (PI) for banana versus wine at the three-hour endpoint (Figure 4F and Figure 7E) was defined as:

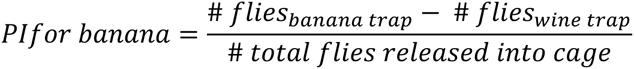

Thus, a preference index of ‘1’ describes a situation in which all 50 flies entered the banana trap, whereas a preference index of ‘-1’ indicates that all 50 flies entered the wine trap. The preference index was computed for each experiment, and then averaged across replicates for a given condition. *Trajectories.* 2D walking trajectories of flies on the surface of the trap platforms were reconstructed post-acquisition from top-camera video data using a custom modification of the Multi tracker software package for tracking multiple objects in 2D^62^, which is built on the ROS (Robot Operating System) framework (https://github.com/tom-f-oconnell/multi_tracker). In brief, Multi tracker performs background subtraction and thresholding, contour identification, splits contours larger than a user-specified size into smaller contours (when two flies touch one another), and applies Kalman filtering to smooth position information and estimate association of data from adjacent frames. Multi tracker extracts trajectory identification numbers, position (*x, y*) in a given frame, and instantaneous velocity (*x, y*). The quality of tracking was manually evaluated by overlaying reconstructed trajectories on minimum intensity projections of the video. If necessary, manual adjustment to tracking settings were made to improve tracking accuracy in each video, which occasionally had slight differences in lighting. For a subset of experiments, individual fly trajectories were proofread by manual comparison to fly video data to confirm the accuracy of trajectory reconstruction.

*Walking speed.* Walking speeds (Figure S2A) were computed from instantaneous (*x, y*) trajectory velocities. For construction of histograms (Figure S2C), walking speeds were averaged over 5-min bins. To evaluate if walking speed changed as a function of how long a fly had been walking on the trap platform (for a given visit to the platform) (Figure S2E), and whether this changes with odor exposure, walking speeds were computed from instantaneous (*x, y*) trajectory velocities grouped in 5-s non-overlapping time bins.

*Occupancy.* The number of fly landings (determined as the initiation of a new trajectory) or the number of flies (determined as the number of unique trajectories) on a trap platform over time was computed in 5-min time bins with 50% overlap, for a total of 72 overlapping bins tiling the three-hour experiment, summed over ten replicates in each condition (Figure 3C and E). Cumulative landing curves were computed by summing landings at each time point across the combined data of the ten experimental replicates for each condition (Figure S3B). For the occupancy analysis in Figure 3E, trajectories were classified as “in ROI” (see below) if they visited at least one entrance hole during the 5-min time bin, and as “out of ROI” if they did not visit any entrance hole during the 5-min time bin. The normalized spatial distribution of flies on the trap platform over time was visualized by performing a 2D kernel density estimation, scaled to a maximum of one for each condition (Figure S3D).

*Approaches.* The region of interest (ROI) corresponding to the region around the entrance holes of each trap was generated by extracting the center coordinate of each entrance hole (using ImageJ) and defining a four-pixel (∼1 mm) radius around each coordinate. An “approach” was defined as a trajectory that touched or entered any of the (noncontiguous) pixels of the ROI. The time of approach was defined as the time when the trajectory first contacts the ROI. The number of approahes over time was computed in 5-min time bins with 50% overlap, summed over ten replicates in each condition (Figure 3G). The number of approaches during the first hour of the experiment was determined for each experiment, without binning, and averaged over ten replicates (Figure 3F). The approach duration (Figure S3F) was defined as the time between when the trajectory first comes in contact with the ROI, and the time when the trajectory leaves the ROI or ends inside the ROI.

#### Exit assay

Exits from the odorized environment were detected using the same procedures as for detecting entries into the standard traps (see above). Flies that never exited the chamber were assigned a dwell time of 3 hours. The spatial occupancy in the exit assay was analyzed from overhead video data at four timepoints – 0.5, 1, 2, and 3 hours – from the initiation of the experiment (designated as the time the odor source is introduced). Since flies were recorded as they moved in three spatial dimensions within the cylindrical chamber, Multi tracker did not perform well on this dataset. Instead, video data was background subtracted, thresholded, and the ImageJ plugin Particle Tracker^63^ was applied to obtain the centroid coordinates for each particle (fly) inside the exit assay. The positions of flies along the long axis of the chamber (*x*-coordinate) extracted by the Particle Tracker were manually reviewed and corrected when necessary at each of the four time points. The coordinates were normalized by dividing the *x*-coordinate value by the total interior length of the chamber interior, which varied very slightly between experiments due to slight differences in the cotton barrier. Histograms constructed from the normalized positions were pooled across the ten replicates for each condition. A value of zero means that the flies are at the nearest position to the odor source as possible, whereas a value of one indicates that the flies are at the farthest position from the odor source (adjacent to the exit).

#### Survival experiments

Videos were loaded into ImageJ, and dead flies were marked with the point tool at one-hour intervals. Dead flies were easily identified; they lay on their sides or backs, did not move in later frames, and changed position only when pushed by other flies. The percentage of dead flies at each time point was averaged across three replicates for each condition.

#### Calcium imaging

Calcium imaging data was analyzed using custom scripts in MATLAB (version R2020b; Mathworks), and final plots were generated in RStudio. Movies of raw fluorescence comprising the three trials for each odor stimulus were motion-corrected (rigid), and a mean projection of each movie was created. For each experiment, the mean projection for one movie (one stimulus) was selected as a template, and the remaining mean projections were aligned to the template using rigid image registration. The vector for image alignment was then applied back onto the whole movies to achieve image registration across all movies of a fly. A total average image across all mean projections was computed and was used for manual creation of a region of interest (ROI) in each fly, which circumscribed all labeled PN axon terminals in the mushroom body calyx in that imaging plane as defined by GCaMP6f fluorescence. ΔF/F videos were created by computing the relative change in fluorescence (ΔF) normalized to the mean fluorescence (F) in the baseline epoch (first 40 frames or ∼8.9 s of the pre-stimulus baseline period), in each pixel of each frame. Change in fluorescence over time in the ROI was determined by computing the mean ΔF/F value across pixels inside the ROI for each frame of the movie, averaged across trials. A seven-frame window starting at the time of nominal stimulus onset was defined. The frame with the maximum ΔF/F in the ROI was designated the peak response for that trial. Heatmaps of peak responses were generated by averaging across the three trials of the stimulus.

*Dose response curves*. The response curves of peak ΔF/F as a function of stimulus concentration were fit to a three-parameter logistic function:

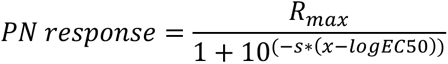

Fits and determination of 95% confidence intervals of model parameters (e.g., EC50) were performed in MATLAB using nlinfit.

*Principal component analysis (PCA)*. Inputs to PCA were the 57 peak response patterns recorded in an individual fly to 19 different odor stimuli. Images were smoothed with a 2D Gaussian kernel with SD 2 prior to PCA. Inputs were centered but not scaled. Pixels outside the ROI were assigned a value of zero.

#### Statistical analyses

Error bars in figures are bootstrapped 95% confidence intervals, unless otherwise indicated.

*Permutation testing.* Two-way statistical comparisons were performed using permutation testing; the null hypothesis was that all samples from the two comparison groups belong to the same distribution. Observations across experiments for the two conditions being compared were combined and randomly reassigned (permuted) into two groups, maintaining the number of observations in each comparison group. The difference between the means of the shuffled groups was calculated. The permutation process was repeated for a total of 10,000 resamplings, and the two-tailed *p*-value for the comparison was computed as the proportion of resamplings in which the absolute difference of the resampled means was larger than the absolute value of the observed difference between experimental groups. Statistical significance reported in all figures reflect Bonferroni-adjustment of *p-*values to correct for multiple comparisons in a given analysis.

For determination of the 95% confidence interval of cumulative entry/exit curves, data across experimental replicates were combined in each condition and resampled (with replacement) at each time point. All flies in the assay, including those that did not enter (or exit) any trap during the three-hour assay period, were included in the analysis. The 0.025 and 0.975 quantile of the 10,000 resampled curves at each time point were taken as the 95% confidence interval. Only flies that landed were included in the calculation of the 95% confidence interval of the cumulative landing curves; since we did not have a quantitative count of the number of flies that did not land they were not included in the analysis.

Comparison of the distributions of times of first entry into a trap between odor exposure groups (Figure 2E, 4C) was performed using a pairwise log-rank test (survival:: pairwise_survdiff, R). Comparisons of survival curves between odor exposure groups or between genotypes (Figure 2B, S3A), and comparisons of empirical cumulative distribution functions (Figure 4G-H), were performed using a two-sample Kolmogorov-Smirnov test (stats::ks.test, R). The non-parametric Spearman’s rank correlation (stats::cor.test, R) was used to measure the monotonic association between the mean number of trap entries and the mean PN output response strength, across odor stimuli in the panel (Figure 1H).

## SUPPLEMENTARY MATERIALS

**Supplementary Figure S1:**
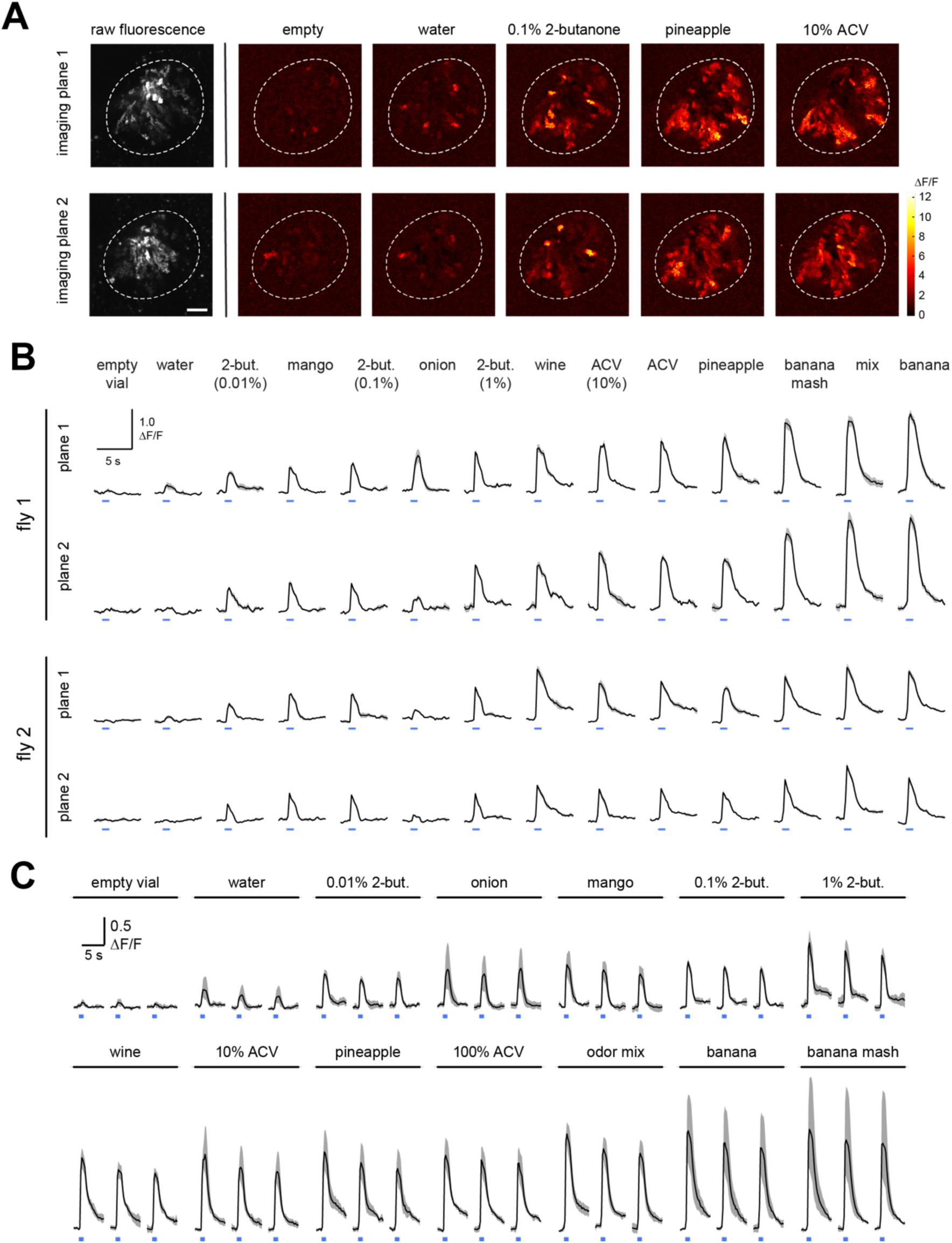
The relative amplitudes of odor-evoked responses in PN axon terminals across odor stimuli is not strongly sensitive to the specific imaging plane in the calyx. Related to Figure 1. **(A)** Peak ΔF/F odor-evoked response patterns in PN axon terminals in the mushroom body calyx (white dashed line) in two different z-planes in an example fly. Grayscale image (left) shows raw fluorescence. Imaging plane 1 (top row) was 6 µm deeper in the calyx than imaging plane 2 (bottom row). ACV, apple cider vinegar. Scale bar, 10 µm. The fly genotype was yw/+; *NP225-Gal4*/*20x-UAS-IVS-Syn21-OpGCaMP6f-p10*; +. **(B)** Time course of odor-evoked changes in fluorescence (mean and S.E.M. across trials) in PN axon terminals in two different imaging planes in the mushroom body calyx for two example flies (three trials/stimulus). Fly 1 is from **(A).** Blue bar indicates the 1-s odor stimulus. Responses to different stimuli are ordered according to their amplitudes in plane 1 of fly 1. Odor stimuli from left to right: empty odor vial; water; 2-butanone (10^-4^); mango; 2-butanone (10^-3^); onion; 2-butanone (10^-2^); red wine; 10% ACV; 100% ACV; pineapple; mashed banana; mix of 2-butanone, 2-pentanone, E2-hexenal, and pentyl acetate, each at 1%; and banana. The order of stimulus presentation was randomized in each fly, but the presentation order was the same for different imaging planes in the same fly to allow a direct comparison. Imaging plane 1 was located ventrally to plane 2 in each calyx. **(C)** Time course of odor-evoked changes in fluorescence (mean and 95% CI) in PN terminals, measured in a single plane of the MB calyx in each fly (n = 5 flies). ΔF/F response was computed as the pixel mean within a large ROI that circumscribed PN axonal boutons in each fly. Blue bar indicates the 1-s odor stimulus. Odor stimuli are as in **(B)**.

**Supplementary Figure S2:**
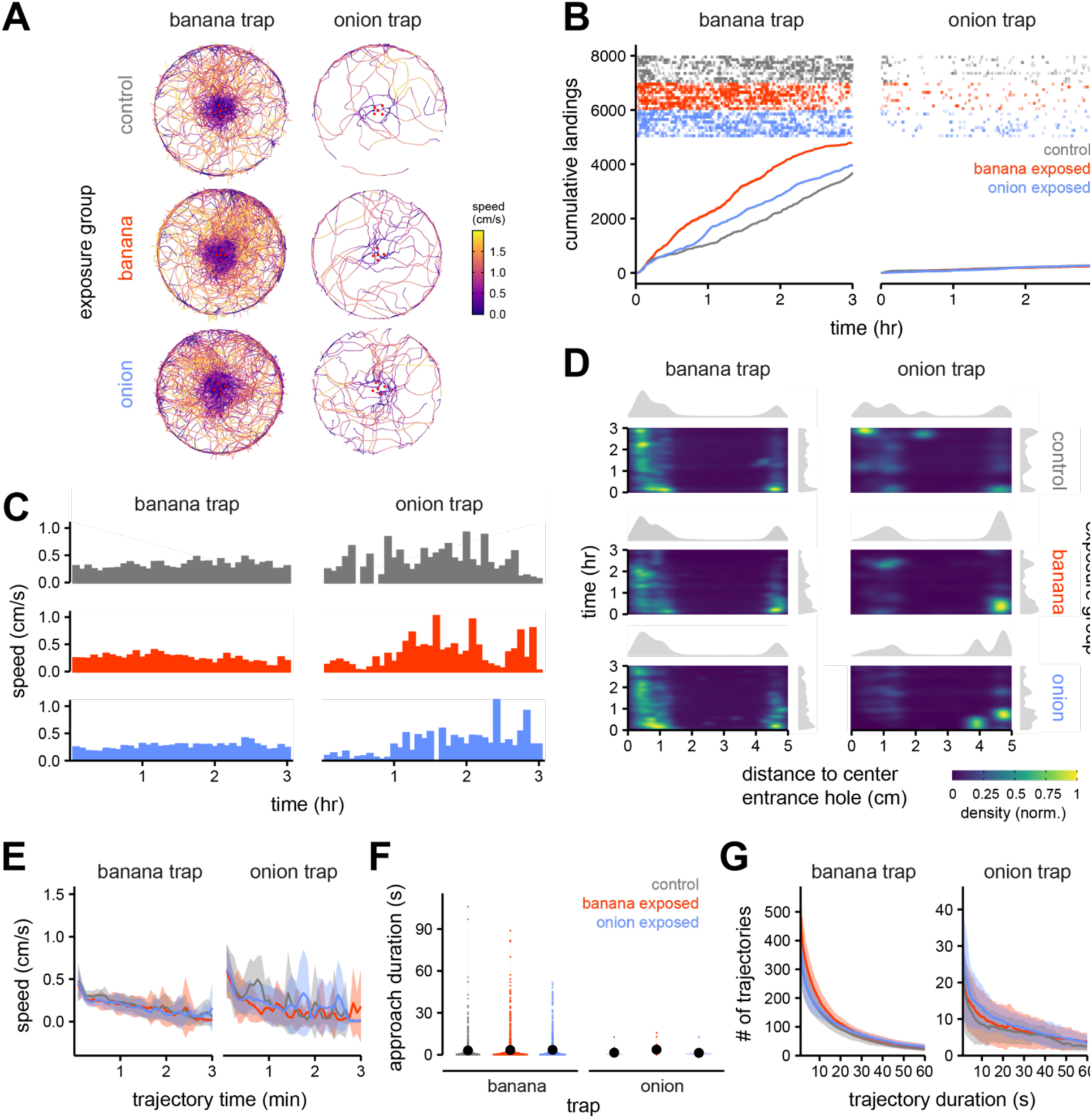
Additional behavioral metrics of flies allowed to choose between a trap baited with onion or banana in the free-flight trap assay. Related to Figures 2 and 3. **(A)** Overlay of all trajectories on each trap platform, color coded by instantaneous walking speed, from an example experiment for each odor exposure condition. The solid red circles are the trap entrances. Color scale is truncated at 2 cm/s. **(B-G)** Quantification of fly behavior in experiments from Figure 2C-E. Control- (grey), banana odor- (red), or onion odor- (blue) exposed flies in free-flight chose between a trap baited with onion or a trap baited with banana (n=10 experiments/condition). **(B)** Landings on each trap for control- (gray), banana odor- (red), or onion odor- (blue) exposed flies. Top rasters: individual landing events over time; each row is an experiment. Bottom: Cumulative landings on each trap across all experiments (n=10) for each odor exposure group. Error envelope is 95% CI and barely visible at this scale. **(C)** Mean walking speed, computed over 5-min bins, as a function of time in the experiment (n = 10 experiments). **(D)** Normalized radial distribution of flies on the trap platform as a function of time in the experiment (mean of 10 experiments). Distance (x-axis) represents the distance from the fly to the center trap entrance (0 cm, center entrance hole; ∼0.5 cm, outer entrance holes; ∼1.5 cm, boundary between odor vial cap and platform disc; ∼5 cm: outer edge of platform. **(E)** Walking speed (mean and SD) as a function of time from the initiation of the trajectory, computed across all trajectories in each condition. The time-axis was truncated at three minutes (mean and SD). **(F)** Distribution of time durations of trajectories that enter the trap entrance ROI (e.g., approaches) for each condition (see Figure 3A). Solid black circles are the means of each condition. Two approaches lasted longer than 250 seconds and were omitted for better display scaling. **(G)** Distribution of trajectories (mean and SD across 10 experiments) across trajectory durations in time. The x-axis is truncated at sixty seconds.

**Supplementary Figure S3:**
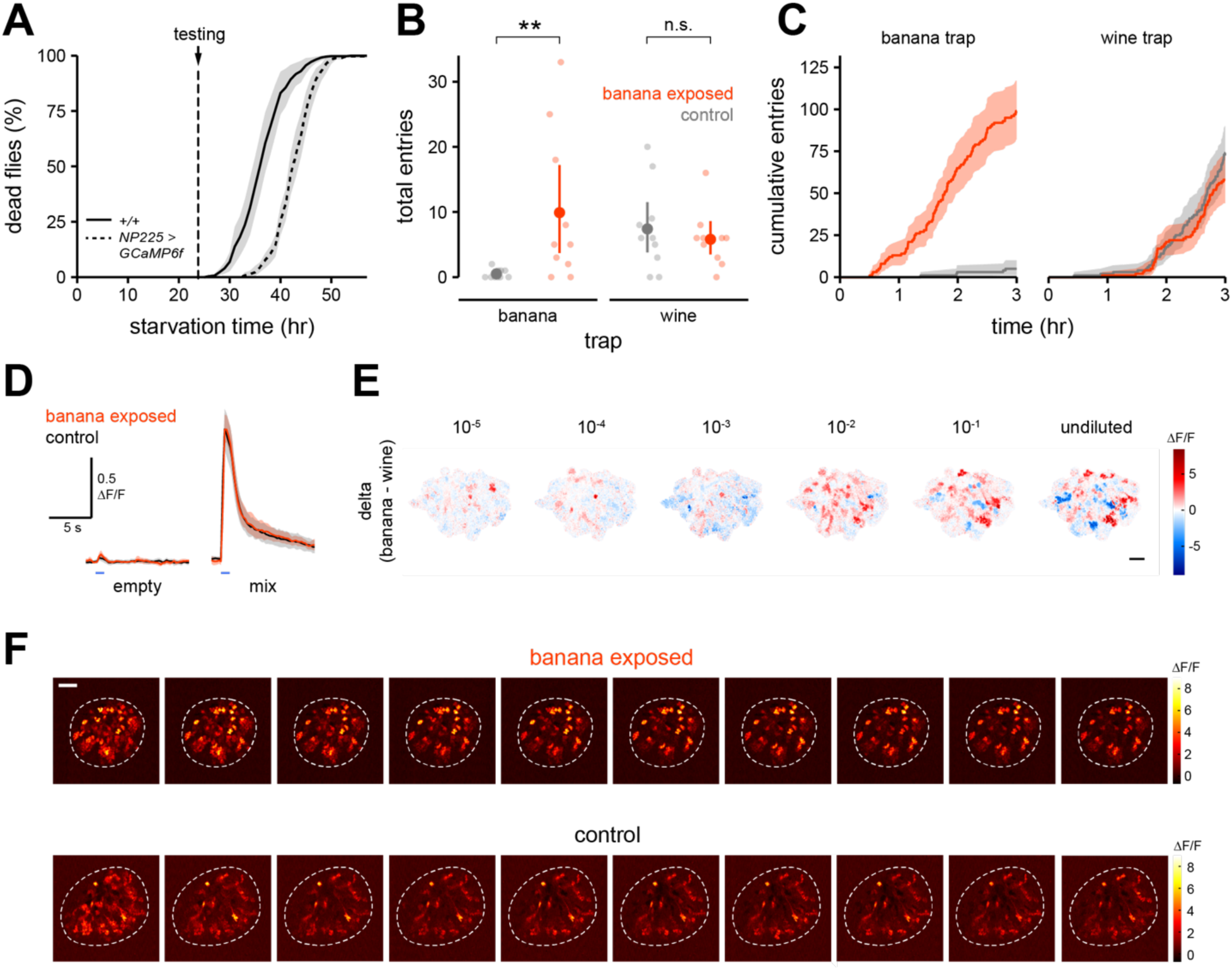
Measurement of odor representations in the PN axon terminals of flies chronically exposed to natural odor. Related to Figure 6. **(A)** Survival curves for wildtype (HCS) (solid line) and *NP225>GCaMP6f* flies (dashed line). Data are mean and 95% CI, averaged across experiments using each genotype exposed to different odors (n = 3 experiments/condition; wildtype: control-, banana odor-, onion odor-exposed; *NP225>GCaMP6f*: control-, banana odor-exposed). Odor exposure was as in Figure 2A; starvation time 0 corresponds to the start of the third day in Figure 2A. Testing would normally commence at 24 hrs starvation time for wildtype flies. The full genotype of … is yw/+; *NP225- Gal4*/*20x-UAS-IVS-Syn21-OpGCaMP6f-p10*; +. **(B)** Total entries (mean and 95% CI, solid symbols) into banana- or wine-odorized traps over three hours by control- (grey) and banana odor- (red) exposed *NP225>GCaMP6f* flies in the free flight trap assay (n = 10 experiments, light symbols). Flies were starved for 30.5 hrs and were assayed in groups of 70 flies/experiment. \*\**p* < 0.01, n.s., no significant difference (*p* ≥ 0.05), two-tailed permutation t-test with Bonferroni correction. **(C)** Cumulative entries into each trap across all experiments for each odor exposure group from **(B)**. Error envelope is the 95% CI bootstrapped across experiments (n = 10). **(D)** Change in fluorescence over time (mean and 95% CI) in PN axon terminals in response to an empty vial stimulus or a mix of 2-butanone, 2-pentanone, E2-hexenal, and pentyl acetate (each at 1%) in banana odor- (red) and control- (gray) exposed *NP225>GCaMP6f* flies (n = 5–11 flies, banana odor-exposed; n = 5—9 flies, control-exposed). Blue bar indicates time of 1s odor pulse. **(E)** Difference between peak ΔF/F response patterns in PN terminals to banana and wine odors at increasing stimulus intensities in a control-exposed fly. Red designates pixels activated more by banana odor than wine odor; blue designates the reverse. Scale bar, 10 µm. **(F)** Peak ΔF/F response images in PN terminals extracted from each successive 1 s pulse in the “fluctuating” banana odor-stimulus (Figure 6F) in an example banana odor-exposed (top row) or control-exposed fly (bottom row). Images are ordered in time, with the response to the first pulse on the left. Scale bar, 10 µm.

**Supplementary Video S1:**
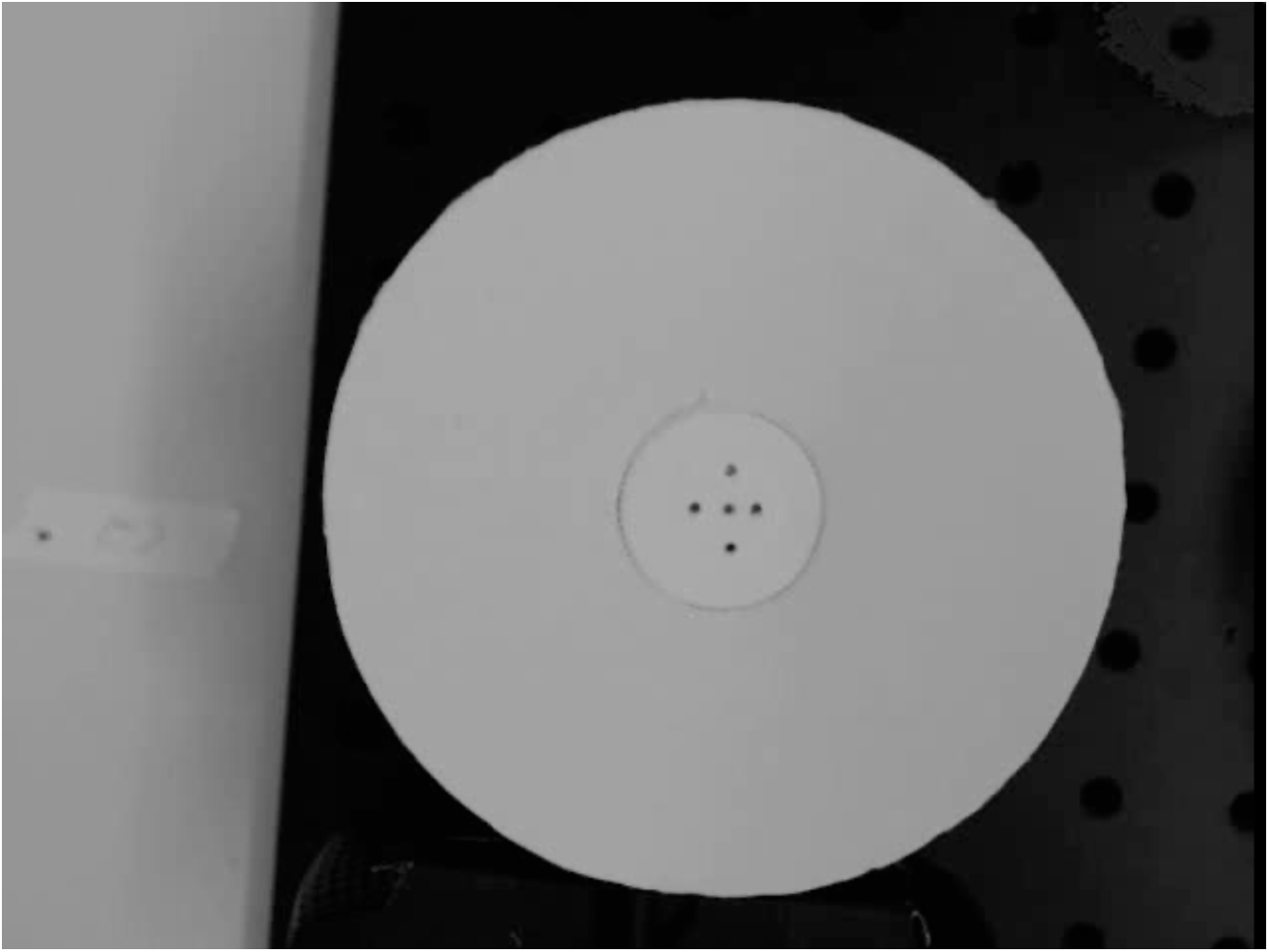
Video from the top-view camera (see Figure 1A) of control odor-exposed flies walking on a trap baited with banana. Multiple approaches to the trap entrances are made by the first fly; later, a second fly on the trap exhibits a similar behavior. Frame rate, 5 Hz.

## REFERENCES

1. Sengpiel, F., Stawinski, P., and Bonhoeffer, T. (1999). Influence of experience on orientation maps in cat visual cortex. Nat Neurosci 2, 727–732. 10.1038/11192.

2. Blakemore, C., and Cooper, G.F. (1970). Development of the brain depends on the visual environment. Nature 228, 477–478.

3. Blasdel, G.G., Mitchell, D.E., Muir, D.W., and Pettigrew, J.D. (1977). A physiological and behavioural study in cats of the effect of early visual experience with contours of a single orientation. The Journal of Physiology 265, 615–636. 10.1113/jphysiol.1977.sp011734.

4. Kreile, A.K., Bonhoeffer, T., and Hübener, M. (2011). Altered Visual Experience Induces Instructive Changes of Orientation Preference in Mouse Visual Cortex. J Neurosci 31, 13911– 13920. 10.1523/JNEUROSCI.2143-11.2011.

5. Zhang, L.I., Bao, S., and Merzenich, M.M. (2001). Persistent and specific influences of early acoustic environments on primary auditory cortex. Nat Neurosci 4, 1123–1130. 10.1038/nn745.

6. Stanton, S.G., and Harrison, R.V. (1996). Abnormal cochleotopic organization in the auditory cortex of cats reared in a frequency augmented environment. Aud. Neurosci 2, 97–107.

7. Ayabe-Kanamura, S., Schicker, I., Laska, M., Hudson, R., Distel, H., Kobayakawa, T., and Saito, S. (1998). Differences in perception of everyday odors: A Japanese-German cross-cultural study. Chemical Senses 23, 31–38. 10.1093/chemse/23.1.31.

8. Poncelet, J., Rinck, F., Bourgeat, F., Schaal, B., Rouby, C., Bensafi, M., and Hummel, T. (2010). The effect of early experience on odor perception in humans: Psychological and physiological correlates. Behavioural Brain Research 208, 458–465. 10.1016/j.bbr.2009.12.011.

9. Pangborn, R.M., Guinard, J.-X., and Davis, R.G. (1988). Regional aroma preferences. Food Quality and Preference 1, 11–19. 10.1016/0950-3293(88)90003-1.

10. Schaal, B., Marlier, L., and Soussignan, R. (2000). Human Foetuses Learn Odours from their Pregnant Mother’s Diet. Chemical Senses 25, 729–737. 10.1093/chemse/25.6.729.

11. Arshamian, A., Gerkin, R.C., Kruspe, N., Wnuk, E., Floyd, S., O’Meara, C., Garrido Rodriguez, G., Lundström, J.N., Mainland, J.D., and Majid, A. (2022). The perception of odor pleasantness is shared across cultures. Current Biology 32, 2061–2066.e3. 10.1016/j.cub.2022.02.062.

12. Mandairon, N., and Linster, C. (2009). Odor Perception and Olfactory Bulb Plasticity in Adult Mammals. Journal of Neurophysiology 101, 2204–2209. 10.1152/jn.00076.2009.

13. Golovin, R.M., and Broadie, K. (2016). Developmental experience-dependent plasticity in the first synapse of the Drosophila olfactory circuit. J Neurophysiol 116, 2730–2738. 10.1152/jn.00616.2016.

14. Wark, B., Lundstrom, B.N., and Fairhall, A. (2007). Sensory adaptation. Curr Opin Neurobiol 17, 423–429. 10.1016/j.conb.2007.07.001.

15. Barlow, H.G. (1961). Possible principles underlying the transformation of sensory messages. In Sensory Communication, W. A. Rosenblith, ed. (MIT Press), pp. 217–234.

16. Laughlin, S. (1981). A simple coding procedure enhances a neuron’s information capacity. Zeitschrift fur Naturforschung. Section C. Biosciences 36, 910–912.

17. Das, S., Sadanandappa, M.K., Dervan, A., Larkin, A., Lee, J.A., Sudhakaran, I.P., Priya, R., Heidari, R., Holohan, E.E., Pimentel, A., et al. (2011). Plasticity of local GABAergic interneurons drives olfactory habituation. Proc Natl Acad Sci U S A 108, E646–E654. 10.1073/pnas.1106411108.

18. Sachse, S., Rueckert, E., Keller, A., Okada, R., Tanaka, N.K., Ito, K., and Vosshall, L.B. (2007). Activity-dependent plasticity in an olfactory circuit. Neuron 56, 838–850. 10.1016/j.neuron.2007.10.035.

19. Han, Y.K., Köver, H., Insanally, M.N., Semerdjian, J.H., and Bao, S. (2007). Early experience impairs perceptual discrimination. Nat Neurosci 10, 1191–1197. 10.1038/nn1941.

20. Muir, D.W., and Mitchell, D.E. (1975). Behavioral deficits in cats following early selected visual exposure to contours of a single orientation. Brain Research 85, 459–477. 10.1016/0006-8993(75)90820-3.

21. Olsen, S.R., and Wilson, R.I. (2008). Cracking neural circuits in a tiny brain: new approaches for understanding the neural circuitry of Drosophila. Trends Neurosci 31, 512–520.

22. Vosshall, L.B., and Stocker, R.F. (2007). Molecular architecture of smell and taste in Drosophila. Annu Rev Neurosci 30, 505–533. 10.1146/annurev.neuro.30.051606.094306.

23. Devaud, J.M., Acebes, A., and Ferrus, A. (2001). Odor exposure causes central adaptation and morphological changes in selected olfactory glomeruli in Drosophila. The Journal of neuroscience : the official journal of the Society for Neuroscience 21, 6274–6282.

24. Devaud, J.M., Acebes, A., Ramaswami, M., and Ferrus, A. (2003). Structural and functional changes in the olfactory pathway of adult Drosophila take place at a critical age. Journal of neurobiology 56, 13–23. 10.1002/neu.10215.

25. Bruce, T.J.A., and Pickett, J.A. (2011). Perception of plant volatile blends by herbivorous insects – Finding the right mix. Phytochemistry 72, 1605–1611. 10.1016/j.phytochem.2011.04.011.

26. Wright, G.A., Lutmerding, A., Dudareva, N., and Smith, B.H. (2005). Intensity and the ratios of compounds in the scent of snapdragon flowers affect scent discrimination by honeybees (apis mellifera). Journal of Comparative Physiology A 191, 105–114.

27. Lahondère, C., Vinauger, C., Okubo, R.P., Wolff, G.H., Chan, J.K., Akbari, O.S., and Riffell, J.A. (2020). The olfactory basis of orchid pollination by mosquitoes. Proceedings of the National Academy of Sciences 117, 708–716.

28. Brat, P., Yahia, A., Chillet, M., Bugaud, C., Bakry, F., Reynes, M., and Brillouet, J.-M. (2004). Influence of cultivar, growth altitude and maturity stage on banana volatile compound composition. Fruits 59, 75–82.

29. Montero-Calderón, M., Rojas-Graü, M.A., and Martín-Belloso, O. (2010). Aroma profile and volatiles odor activity along gold cultivar pineapple flesh. Journal of Food Science 75, S506– S512.

30. Pino, J.A., and Febles, Y. (2013). Odour-active compounds in banana fruit cv. Giant Cavendish. Food Chemistry 141, 795–801. 10.1016/j.foodchem.2013.03.064.

31. Becher, P.G., Flick, G., Rozpędowska, E., Schmidt, A., Hagman, A., Lebreton, S., Larsson, M.C., Hansson, B.S., Piškur, J., and Witzgall, P. (2012). Yeast, not fruit volatiles mediate D rosophila melanogaster attraction, oviposition and development. Functional Ecology 26, 822– 828.

32. Budick, S.A., and Dickinson, M.H. (2006). Free-flight responses of Drosophila melanogaster to attractive odors. Journal of Experimental Biology 209, 3001–3017.

33. Faucher, C.P., Hilker, M., and Bruyne, M. de (2013). Interactions of Carbon Dioxide and Food Odours in Drosophila: Olfactory Hedonics and Sensory Neuron Properties. PLOS ONE 8, e56361. 10.1371/journal.pone.0056361.

34. Barrows, W.M. (1907). The reactions of the Pomace fly, Drosophila ampelophila loew, to odorous substances. Journal of Experimental Zoology 4, 515–537. 10.1002/jez.1400040403.

35. Landolt, P., Adams, T., and Rogg, H. (2012). Trapping spotted wing drosophila, Drosophila suzukii (Matsumura)(Diptera: Drosophilidae), with combinations of vinegar and wine, and acetic acid and ethanol. Journal of applied entomology 136, 148–154.

36. Tanaka, N.K., Endo, K., and Ito, K. (2012). Organization of antennal lobe-associated neurons in adult Drosophila melanogaster brain. Journal of Comparative Neurology 520, 4067– 4130. 10.1002/cne.23142.

37. Brenner, N., Bialek, W., and Van Steveninck, R. de R. (2000). Adaptive rescaling maximizes information transmission. Neuron 26, 695–702.

38. Pech, U., Revelo, N.H., Seitz, K.J., Rizzoli, S.O., and Fiala, A. (2015). Optical Dissection of Experience-Dependent Pre- and Postsynaptic Plasticity in the Drosophila Brain. Cell Reports 10, 2083–2095. 10.1016/j.celrep.2015.02.065.

39. Chen, T.W., Wardill, T.J., Sun, Y., Pulver, S.R., Renninger, S.L., Baohan, A., Schreiter, E.R., Kerr, R.A., Orger, M.B., Jayaraman, V., et al. (2013). Ultrasensitive fluorescent proteins for imaging neuronal activity. Nature 499, 295–300. 10.1038/nature12354.

40. Kidd, S., Struhl, G., and Lieber, T. (2015). Notch is required in adult Drosophila sensory neurons for morphological and functional plasticity of the olfactory circuit. PLoS genetics 11, e1005244. 10.1371/journal.pgen.1005244.

41. Chakraborty, T.S., Goswami, S.P., and Siddiqi, O. (2009). Sensory Correlates of Imaginal Conditioning in Drosophila melanogaster. Journal of Neurogenetics 23, 210–219. 10.1080/01677060802491559.

42. Mandairon, N., Stack, C., Kiselycznyk, C., and Linster, C. (2006). Broad activation of the olfactory bulb produces long-lasting changes in odor perception. PNAS 103, 13543–13548. 10.1073/pnas.0602750103.

43. Wang, H.W., Wysocki, C.J., and Gold, G.H. (1993). Induction of olfactory receptor sensitivity in mice. Science 260, 998–1000. 10.1126/science.8493539.

44. Cadiou, H., Aoudé, I., Tazir, B., Molinas, A., Fenech, C., Meunier, N., and Grosmaitre, X. (2014). Postnatal Odorant Exposure Induces Peripheral Olfactory Plasticity at the Cellular Level. J. Neurosci. 34, 4857–4870. 10.1523/JNEUROSCI.0688-13.2014.

45. Liu, A., and Urban, N.N. (2017). Prenatal and Early Postnatal Odorant Exposure Heightens Odor-Evoked Mitral Cell Responses in the Mouse Olfactory Bulb. eNeuro 4. 10.1523/ENEURO.0129-17.2017.

46. Chodankar, A., Sadanandappa, M.K., VijayRaghavan, K., and Ramaswami, M. (2020). Glomerulus-Selective Regulation of a Critical Period for Interneuron Plasticity in the Drosophila Antennal Lobe. J. Neurosci. 40, 5549–5560. 10.1523/JNEUROSCI.2192-19.2020.

47. Dunkel, A., Steinhaus, M., Kotthoff, M., Nowak, B., Krautwurst, D., Schieberle, P., and Hofmann, T. (2014). Nature’s chemical signatures in human olfaction: a foodborne perspective for future biotechnology. Angewandte Chemie International Edition 53, 7124–7143.

48. Maarse, H. ed. (2017). Volatile Compounds in Foods and Beverages (Routledge)10.1201/9780203734285.

49. Nijssen L. M. (1996). Volatile Compounds in Food : Qualitative and Quantitative Data 7th ed. (Zeist Netherlands: TNO-CIVO Food Analysis Institute).

50. Jordán, M.J., Tandon, K., Shaw, P.E., and Goodner, K.L. (2001). Aromatic Profile of Aqueous Banana Essence and Banana Fruit by Gas Chromatography−Mass Spectrometry (GC-MS) and Gas Chromatography−Olfactometry (GC-O). J. Agric. Food Chem. 49, 4813– 4817. 10.1021/jf010471k.

51. Gugel, Z.V., Maurais, E., and Hong, E.J. (2021). Chronic exposure to odors at naturally occurring concentrations triggers limited plasticity in early stages of Drosophila olfactory processing. bioRxiv, 10.1101/2021.09.03.458834.

52. National Institute for Occupasional Safety and Health Immediately Dangerous To Life or Health (IDLH) Values. https://www.cdc.gov/niosh/idlh/default.html.

53. Tempel, B.L., Bonini, N., Dawson, D.R., and Quinn, W.G. (1983). Reward learning in normal and mutant Drosophila. Proceedings of the National Academy of Sciences 80, 1482–1486. 10.1073/pnas.80.5.1482.

54. Schwaerzel, M., Monastirioti, M., Scholz, H., Friggi-Grelin, F., Birman, S., and Heisenberg, M. (2003). Dopamine and Octopamine Differentiate between Aversive and Appetitive Olfactory Memories in Drosophila. J. Neurosci. 23, 10495–10502. 10.1523/JNEUROSCI.23-33-10495.2003.

55. Law, C.-T., and Gold, J.I. (2009). Reinforcement learning can account for associative and perceptual learning on a visual-decision task. Nature neuroscience 12, 655–663.

56. Kass, M.D., Moberly, A.H., Rosenthal, M.C., Guang, S.A., and McGann, J.P. (2013). Odor-Specific, Olfactory Marker Protein-Mediated Sparsening of Primary Olfactory Input to the Brain after Odor Exposure. J. Neurosci. 33, 6594–6602. 10.1523/JNEUROSCI.1442-12.2013.

57. Kay, L.M., and Laurent, G. (1999). Odor- and context-dependent modulation of mitral cell activity in behaving rats. Nature Neuroscience 2, 1003–1009.

58. Chu, M.W., Li, W.L., and Komiyama, T. (2016). Balancing the robustness and efficiency of odor representations during learning. Neuron 92, 174–186.

59. McGann, J.P. (2015). Associative learning and sensory neuroplasticity: how does it happen and what is it good for? Learning & Memory 22, 567–576.

60. Hong, E.J., and Wilson, R.I. (2015). Simultaneous encoding of odors by channels with diverse sensitivity to inhibition. Neuron 85, 573–589. 10.1016/j.neuron.2014.12.040.

61. Zocchi, D., Emily, S.Y., Hauser, V., O’Connell, T.F., and Hong, E.J. (2022). Parallel encoding of CO2 in attractive and aversive glomeruli by selective lateral signaling between olfactory afferents. Current Biology.

62. van Breugel, F., Huda, A., and Dickinson, M.H. (2018). Distinct activity-gated pathways mediate attraction and aversion to CO 2 in Drosophila. Nature 564, 420–424. 10.1038/s41586-018-0732-8.

63. Sbalzarini, I.F., and Koumoutsakos, P. (2005). Feature point tracking and trajectory analysis for video imaging in cell biology. Journal of structural biology 151, 182–195.

